# The use of Ipratropium and Tiotropium as novel agents to reduce inflammation in *in vitro* macrophage models

**DOI:** 10.1101/2021.10.31.466274

**Authors:** Ifteqar Hussain Mohammed

## Abstract

**Background:** Chronic Obstructive Pulmonary Disease (COPD) affects an estimated 330 million individuals worldwide. Approximately, 3 million individuals died of COPD in 2012 and it is predicted that COPD would be the third leading factor for deaths worldwide by 2020. In United Kingdom nearly one million individuals suffer from COPD.

**Purpose:** There are no effective pharmacotherapies available for COPD. it is only managed by using bronchodilators and inhaled corticosteroids mostly. However, cardiovascular effects are associated with these drugs. Most importantly, there is an unmet need of COPD treatment worldwide. Our research aim was to identify Ipratropium and Tiotropium as novel anti-inflammatory agents in *in vitro* macrophage models.

**Aims:** To investigate the LPS stimulated pro-inflammatory cytokines IL-6 and TNF- α levels in THP-1 cells. To investigate whether the drugs Ipratropium and Tiotropium are capable of decreasing LPS-induced inflammation in THP-1 cells.

**Materials:** Human monocytic cell line THP-1 cells, Rosewell Park Memorial Institute RPMI 1640 with Glutamax I, 1% Penicillin Streptomycin (PenStrep) and 10% foetal bovine serum (FBS), Lipopolysaccharide 10μl/ml, 0.05% Tween20, 0.4% Trypan blue, Reagent diluent (10% Bovine Serum Albumin in PBS), Budesonide Fenoterol, Ipratropium and Tiotropium. Human IL-6 DuoSet ELISA, Human TNF-α ELISA, TMB ELISA Substrate solution and Stop solution.

**Methods:** THP-1 cells were cultured and challenges with LPS to stimulate the IL-6 and TNF-α cytokines. The cells were treated with Budesonide, Fenoterol, Ipratropium and Tiotropium. ELISA was performed to determine the concentrations of cytokines.

**Results:** The results suggested that Ipratropium and Tiotropium reduce IL-6 and TNF- α concentrations in the cells. However, Budesonide and Fenoterol were found to reduce cytokines more effectively than Ipratropium and Tiotropium. The data was considered significant only when *P* <0.05.

**Conclusions:** The anti-inflammatory or cytokine reducing properties of Ipratropium and Tiotropium were acknowledged. The research hypothesis was found to be true. Budesonide and Fenoterol substantially reduce cytokine levels. The receptor interactions of Ipratropium and Tiotropium may be responsible for their duration of action. Overall, Ipratropium and Tiotropium display the characteristics of novel anti-inflammatories.

## CHAPTER 1 INTRODUCTION

### 1.1 Background of Chronic obstructive pulmonary disease (COPD)

Chronic Obstructive Pulmonary Disease (COPD) can be described as a progressive lung disease which is characterized by the inflammation of lung parenchyma and airway passage. Emphysema, as well as recurrent bronchitis are implicated with COPD (Burney *et al.*, 2003). It is predicted that COPD would be the third leading factor of deaths by 2020 (Pahal, Hashmi and Sharma, 2019). Approximately three million individuals died of COPD in 2012, accounting for six percent of the world’s fatalities (Snell *et al.*, 2016). Over 800,000 patients in England have severe COPD and innumerable number of individuals are undiagnosed (Murphy, 2011). A substantial proportion of the population of the United Kingdom (UK) are affected by COPD. It is the main reason for frequent hospitalization and emergency visits to the healthcare centres. Moreover, it causes serious morbidities such as cardiovascular diseases, diabetes mellitus, dyspnoea and also results in premature deaths (Chen *et al.*, 2015). COPD related fatalities account for nearly 33% of deaths caused by lung disease as shown in figure 1.1. Apparently, COPD is the next most severe pulmonary disease following lung cancer (Hse.gov.uk, 2018).

**Figure 1.1.**
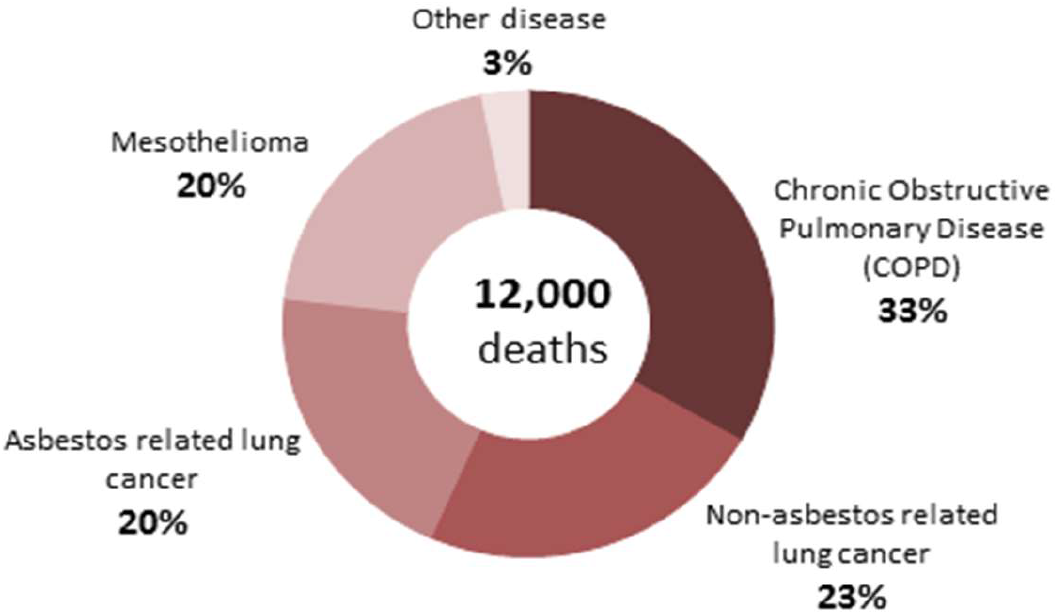
(Hse.gov.uk, 2018) illustrates the various pulmonary diseases contributing to the annual number of deaths in Great Britain.

The National Health Services (NHS) spends approximately £1.9 billion annually for the treatment of COPD. The prevalence of respiratory diseases is higher in patients belonging to socially disadvantaged communities (NHS England, 2019). Such marginalised groups are disadvantaged with a high smoking rate and these groups are more likely to be exposed to occupational dusts, noxious gases, chemicals and environmental pollutants. More than 50% of long-term cigarette smokers are likely to develop COPD. The clinical manifestations usually emerge in individuals after the age of forty but can appear earlier than expected (Chapman *et al.*, 2006).

Several efforts have been made to categorise COPD into distinct groups on clinical and radiological grounds. Nevertheless, it was difficult to diagnose such phenotypes in large population because the disease pathways were not specifically linked (Burgel *et al.*, 2010). Genetics factors are also responsible for development of COPD. The deficiency of serine protease inhibitors such as alpha-1 antitrypsin (AATD) increases emphysema risk (Stoller and Aboussouan, 2005). Genomic studies have suggested that genetic variations in hedgehog proteins and receptors for alpha-nicotinic acetylcholine may influence the susceptibility of COPD (Cho *et al.*, 2014). Nevertheless, the specific role of these gene markers remains uncertain.

Inflammation is the typical immunological response observed when the lungs are exposed to industrial or cigarette smoke. Physiological modifications such as restriction of airways and deterioration of lung parenchyma is observed which not entirely reversible (Soriano *et al.*, 2005).

### 1.2 Background of Asthma

Asthma is a persistent inflammatory disorder of the pulmonary airways. Narrowing of airways in the lungs causes breathlessness, coughing and wheezing. These exacerbations are due to airflow obstruction induced by chronic inflammation in the respiratory system (Boonpiyathad *et al.*, 2019). The pathological feature of asthma is usually airway hyper-responsiveness. However, these conditions are reversible, and it can be treated by anti-inflammatory drugs. Over 300 million individuals worldwide suffer from asthma and it is a serious public health concern. An additional 100 million individuals will be affected by asthma by the year 2025 (Masoli *et al.*, 2004). An estimated 200,000 deaths across the world occur each year due to asthma (Braman, 2006). The incidence rates and severity of asthma in children are high. Whereas, mortality and morbidity are relatively high in adults. Moreover, women over the age of 45-50 are more likely to be affected by asthma (Dharmage, Perret and Custovic, 2019). Nearly 50% of the children in cohort of 20 children diagnosed for respiratory illnesses suffer from severe asthma as shown in figure 1.2 (Anagnostou *et al.*, 2012).

**Figure 1.2.**
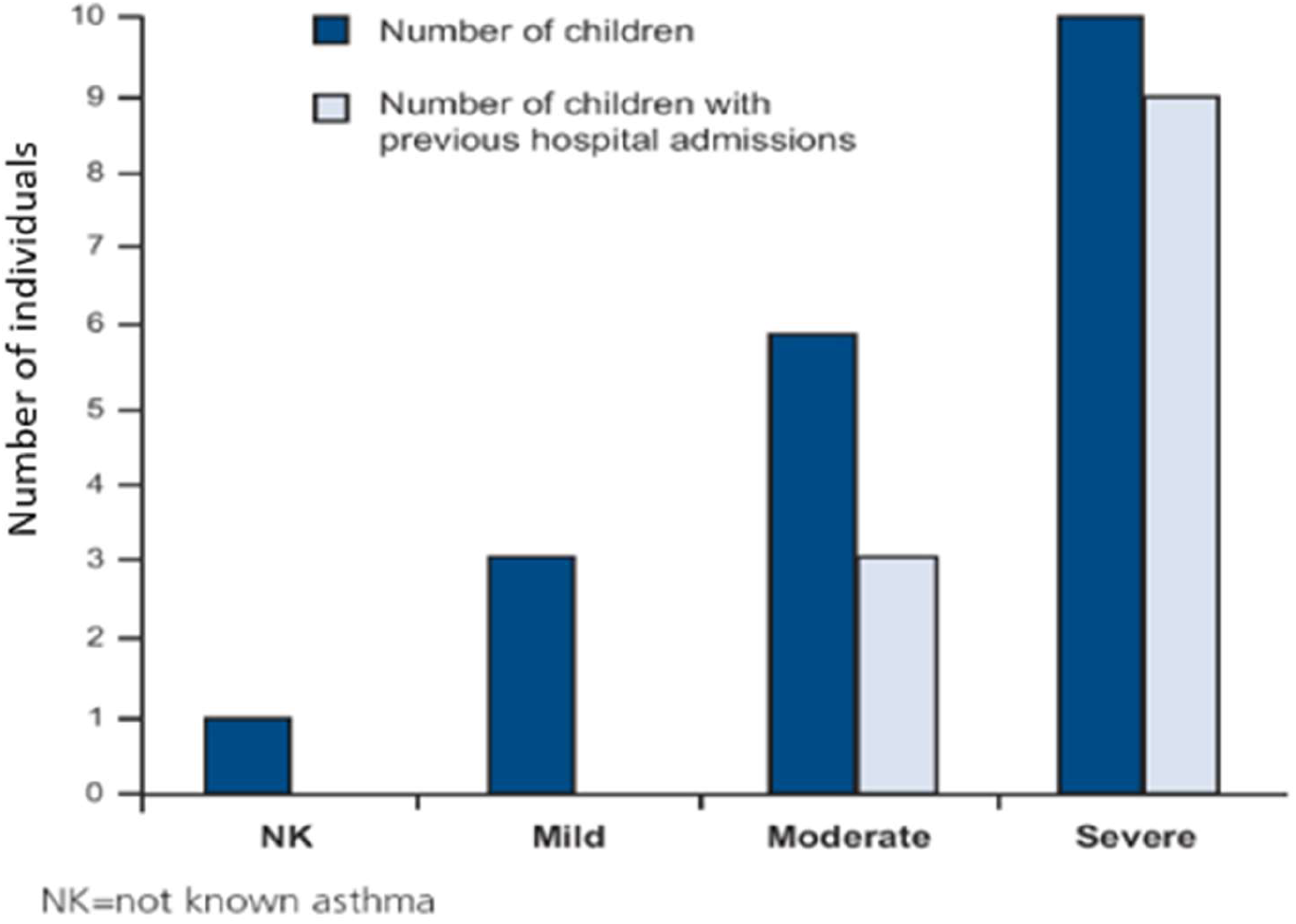
(Anagnostou *et al.*, 2012) illustrates the number of previously and lately hospitalised children in three groups mild, moderate and severe.

The estimated economic burden of Asthma on the NHS is nearly £3 billion (NHS England, 2019). Pharmacotherapy is the most appropriate treatment to control and prevent exacerbations (Ukena, Fishman and Niebling, 2008).

### 1.3 Inflammation

Inflammation in general can be regarded as an immunological response to various pathogens and viruses which enter the human body. Inflammation is an essential defence mechanism which provides protection against foreign particles and maintains homeostasis (Leitch *et al.*, 2008). A number of proinflammatory mediators such as leukocytes and polymorphonuclear granulocytes, agranulocytes, natural killer (NK) cells, mast cells, epithelial cells accumulate at the site of infection to produce immunological response (Robb *et al.*, 2016). Prolonged and uncontrolled anti-inflammatory responses can cause acute or chronic illnesses (Nathan and Ding, 2010). COPD and asthma are the most frequently reported inflammatory diseases. The restriction of airway passage is the common symptom of both diseases, but the underlying inflammatory mechanisms are separate and distinct (Barnes, 2008).

#### 1.3.1 Inflammation in COPD and Asthma

Cytokines are signalling proteins which instigate chronic inflammation in asthma and COPD by recruiting, stimulating and promoting inflammatory cells in the respiratory system. More than 50 cytokines have been recognised which are associated to COPD and asthma related inflammation, but their pathophysiological function is elusive. Cytokines are classified into different categories such as interferons, interleukins, tumour (IL) necrosis factors (TNF), transforming growth factor (TGF) and chemokines (Lordan, Tsoupras and Zabetakis, 2019). The amount of pro-inflammatory cytokines in the bronchoalveolar lavage (BAL) and sputum is significantly high, and these cytokines also amplify exacerbations by activating nuclear factor-kappa B (NF-κB) and p38 mitogen-activated protein kinase signalling pathway which increases the expression of inflammatory genes (Barnes, 2008).

Cigarette smoke upon inhalation triggers the epithelial cells and lung macrophages to release various cytokines. The cytokines such as growth factors TGF-β and FGFs are released which promote the proliferation of fibroblast and result in fibrosis of lung airways as shown in figure 1.3. The cytokines TNF- α, IL-6, IL-1 β and various chemokines (CC) released from the alveolar macrophages amplify the inflammation by attracting numerous monocytes. CCL2 of chemokine family acts via CCR2 and attracts monocytes. The CXCL1 and CXCL8 act through CXCR2 path to attract monocytes and neutrophils. Cytokines belonging to chemokine family CXCL9, CXCL10 and CXCL11 act by Th1 and Tc1 cells and stimulate the release of IFN-γ and results in inflammation. The proteases MMP-9 and neutrophil elastase release EGF and TGF-α which results in mucus hypersecretion as shown in figure 1.3 (Barnes, 2008).

**Figure 1.3.**
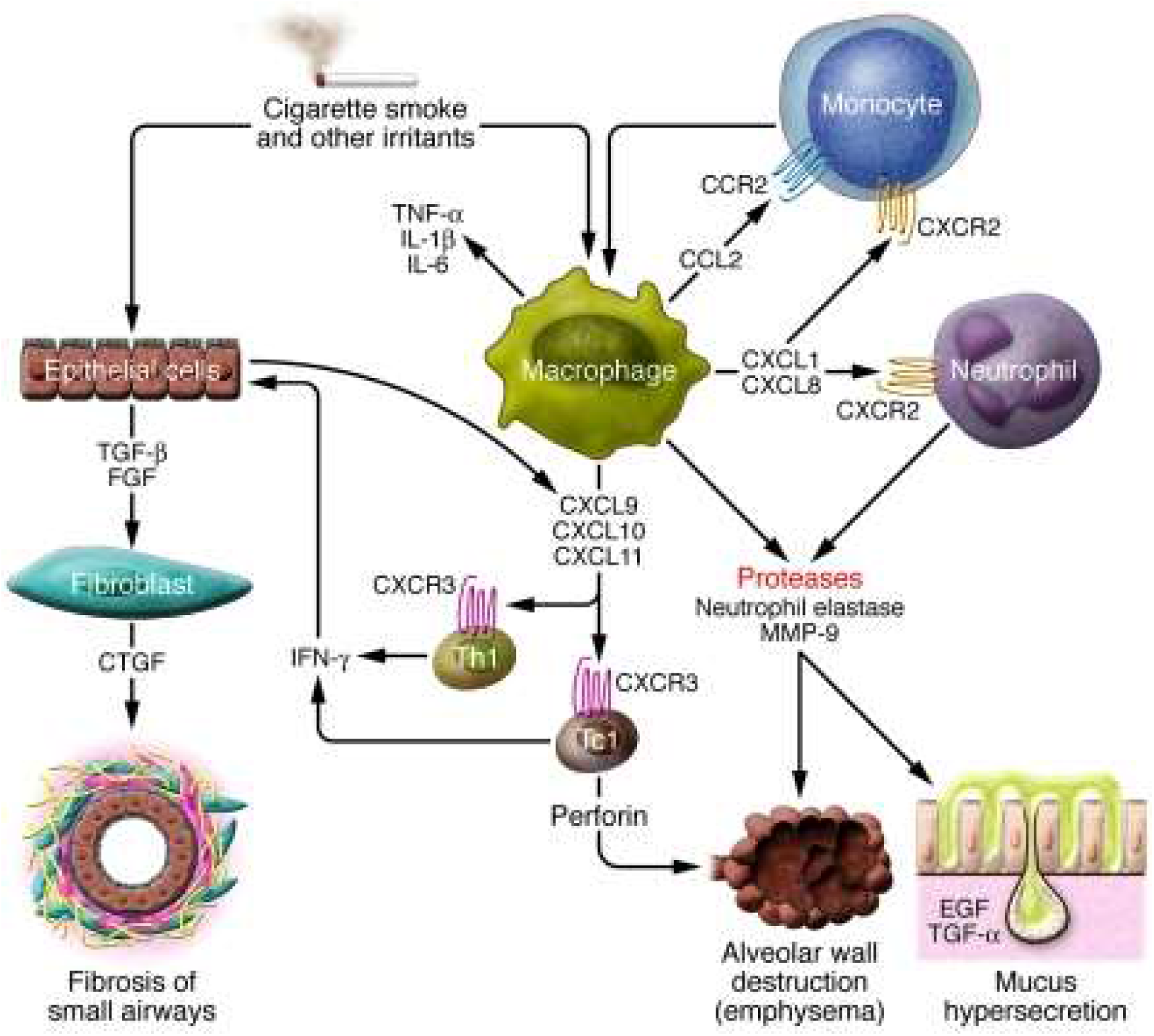
Depicts the cytokines involved in COPD (Barnes, 2008).

The key mediators of inflammation in asthma are stem cell factor (SCF) and thymic stromal lymphoprotein (TSLP) and chemokines CCL5, CCL11, etc. Th2 cells release IL-4 and IL-13 which stimulates B cells to produce immunoglobulin IgE and result in bronchial constriction (Robinson, 2004). Eosinophilic inflammation mediators IL-5 and mast cell proliferators IL-9 are also responsible for inflammation. Prostaglandin D2 and cyteinyl-leukotrines (Cys-LTs) and histamine derived from mast cells results in bronchial constriction and they are also involved in the recruitment of Th2 cells in the airways (Robinson, 2004). The above-mentioned cytokines are the main inflammatory cytokines involved in asthma.

COPD and asthma have similar clinical characteristics but their inflammatory pathways and patterns and response to drug therapy have clear differences (Barnes, 2008). These cytokines are released through various endocrine or paracrine pathways in response to infections affecting immune system by pro-inflammatory mechanisms. Cytokines such as IL-1, IL-6, IL-12, TNF-α and IFN-γ induce inflammation (Ulloa, 2005).

Asthma and COPD can coexist. In both conditions the chronic inflammation of airways is very distinct (Hogg, 2004). However, asthmatic patients when exposed to cigarette smoke and noxious agents may develop airflow limitation and a combination of “asthma-like” or “COPD-like” inflammation (Tillie-Leblond, Gosset and Tonnel, 2005). Furthermore, epidemiological studies suggest that chronic asthma alone may lead to permanent airflow restriction (Bumbacea *et al.*, 2004). Few patients suffering from COPD may exhibit mixed inflammatory symptoms resembling to asthma due to increased eosinophils. While asthma is typically distinguishable from COPD, but it is difficult to differentiate between two diseases in few patients with complicated chronic respiratory problems and fixed airflow obstruction as shown in Table 1 (Cukic *et al.*, 2012).

**Table 1.** (Cukic *et al.*, 2012) shows the main differences in pulmonary inflammation in asthma and COPD.

|  | COPD | ASTHMA | SEVERE ASTHMA |
| --- | --- | --- | --- |
| Cells | Neutrophils ++<br>Macrophages +++<br>CD8+T cells (Tc1)<br>CD4+Tcells (Th2)<br>Th17 | Eosinophils ++<br>Macrophages +<br>Mastocytes<br>CD4+Tcells (Th2) | Neutrophils +<br>Macrophages<br>CD4+Tcells (Th2)<br>CD8+T cells(Tc1)<br>Th17 |
| Key mediators | IL- 8<br>TNFα,IL-1β.IL-6<br>NO+ | Eotaxin<br>IL-4,IL-5,IL-13,<br>NO+++ | IL-8<br>IL-5,IL-13<br>NO++ |
| Oxidative stress | +++ | + | +++ |
| Site of disease | Peripheral airways<br>Lung parenchyma<br>Pulmonary vessels | Proximal airways | Proximal airways<br>Peripheral airways |
| Consequences | Squamous metaplasia<br>Mucous metaplasia,<br>Small airway fibrosis<br>Parenchymal<br>destruction, Pulmonary<br>vascular remodeling | Fragile epithelium<br>Mucous metaplasia,<br>↑ Basement<br>membrane,<br>Bronchoconstriction |  |
| Response to therapy | Small b/d response<br>Poor response to<br>steroids | Large b/d response<br>Good response to<br>steroids | Smaller b/d<br>response<br>Reduced response<br>to steroids |

*NO* = Nitric oxide, *b/d* = bronchodilator.

#### 1.3.2 Effects of inflammation

The structural consequences of inflammation are different in COPD and asthma. Bronchoconstriction occurs in asthma due to the release of inflammatory mediators from the mast cells. The contractility of airway smooth muscles of patients suffering from asthma is usually high and the bronchodilator responses are relatively reduced (Black *et al.*, 2012). Impaired muco-ciliary function leads to obstruction of airways in COPD and asthma. The expression of mucin gene is regulated by Epidermal growth factor receptor (EGFR) via growth factor TGF- α or by neutrophil elastase (NE). This explains the link between chronic bronchitis and neutrophilic inflammation in asthma and COPD patients. (Burgel and Nadel, 2004). Neuronal pathways are responsible for the release of acetylcholine which acts on muscarinic receptors and leads to bronchial constriction. Prolonged fibrosis leads to irreversible bronchoconstriction in COPD (Barnes, 2017).

Cytokines orchestrate the pulmonary inflammation in COPD and asthma. They are potential therapeutic targets for both diseases. Cytokine inhibitors can immensely reduce inflammation by complete or partial blocking of cytokine receptor and also by inhibiting the protein which facilitate signal transduction and transcription via mitogen-activated protein kinase MAPK or NF-κB pathways (Chung, 2006). Grounded on this a range of cytokine inhibitors which can manipulate the immunological network are being studied (Vyas and Vohora, 2016). Blocking of TNF-α cytokine with etanercept (human TNFR-IgG1) improves lung function in patients suffering from asthma (Berry *et al.*, 2006). Patients diagnosed with COPD have increased levels of TNF-α in their sputum. Overall studies suggest that TNF-α inhibitors can be used to treat COPD and asthma. Levels of IL-6 cytokines are significantly high in patients suffering from COPD and asthma (Bhowmik *et al.*, 2000). Pro-inflammatory effect of IL-6 is due to expansion of T helper cells Th2 and Th17 and due to release of C-reactive protein from the hepatic cells. Reduction in IL-6 levels can lead to airway Therefore, treatment of COPD and asthma by inhibition of IL-6 and TNF-α cytokines can be considered as a novel therapeutic approach (Bumbacea *et al.*, 2004).

### 1.4 Treatment of asthma and COPD

The primary objective of asthma treatment is to minimize inflammation and achieve clinical control (Bateman *et al.*, 2008). The goal of COPD pharmacotherapy is to reduce the clinical manifestations, occurrence and severity of exacerbations (Viegi *et al.*, 2007). Almost same medications are used for the treatment of asthma and COPD, but the order and efficiency in treatment is distinct.

Inhalated corticosteroids (ICS) in asthmatic individuals decreases the frequency of manifestations and improves lung function (Hondras, Linde and Jones, 2002). In COPD, ICS are specifically used to prevent frequent exacerbations and to achieve higher bronchodilatory response (Molimard and Colthorpe, 2015). Combined therapy of long-acting β-2 agonists and ICS in treatment of COPD and asthma has immense clinical benefits. In both diseases early diagnosis and treatment can affect morbidity and mortality (Calverley, 2003).

#### 1.4.1 COPD Pharmacotherapy

The main objective of COPD pharmacotherapy is to improve patient health, reduce exacerbations and to maintain lung function.

##### Bronchodilators

Long acting bronchodilators improve inspiration volume and promote expiration of trapped-air in the lungs (Cooper, 2009). The consequent increase in inspiratory volume decreases dyspnoea and improves tolerance to physical exercises (Beeh *et al.*, 2011). β-2 adreno receptor agonists (β-2 agonists) present in smooth muscles binds to β-2 adrenergic receptor. Receptor interaction with G-proteins results in synthesis of cyclic adenosine monophosphate (cAMP) which leads to activation of protein kinase A (Hanania *et al.*, 2002).

Short acting β-2 adrenergic receptor agonist (SABA’s) are used in both acute and chronic conditions. Salbutamol is the most commonly used SABA it acts for 4-6 hours. Patient compliance is relatively high in SABA’s treatment (Appleton *et al.*, 2006). The duration of action of long acting β-2 agonists (LABA’s) is generally more than 12 hours. These drugs improve the forced expiratory volume FEV1 of lungs and exacerbation rate (Tam *et al.*, 2016).

##### Muscarinic Antagonists

The para-sympathetic action of five distinct muscarinic receptors (M_1_-M_5_) mediate the contraction of smooth airway muscles and also the release of mucus in airway lumen. M_1_ receptors are located in peri-bronchial ganglia and M3 receptors are present on bronchial smooth muscles. Acetyl choline release due to physiological conditions results in bronchial constriction; this can be prevented by selectively blocking M_1_ and M_3_ receptors (Lipson, 2006).

Short-acting muscarinic antagonists (SAMAs) are used for treating acute and progressive COPD. Ipratropium bromide is a short-acting muscarinic receptor antagonist and it is used widely for management of COPD. It blocks the M_2_ receptor and also the M_1_ and M_3_ with specific affinity which leads to bronchodilation (Barnes, 2000). Ipratropium inhibits the production of cyclic guanosine monophosphate (cGMP). The concentration of calcium and calcium calmodulin complexes are reduced in the smooth muscles. The onset of action is 1 to 2 hours and the effect last up to 8-12 hours (Lipson, 2006).

Long-acting muscarinic antagonists (LAMAs) such as tiotropium binds to the M_1_ and M_3_ receptors and it dissociates gradually from the receptors due to which it delivers a long-acting effect (Lipson, 2006). Tiotropium binds to M_3_ receptors and prevents the activation of phospholipase C (PLC) and Gq pathways. Consequently, the synthesis of inositol 1,4,5-trisphosphate (IP3) is blocked. As a result, the calcium (Ca2+) mediated bronchoconstriction is prevented as shown in figure 1.5 (Pelaia *et al.*, 2015).

**Figure 1.4.**
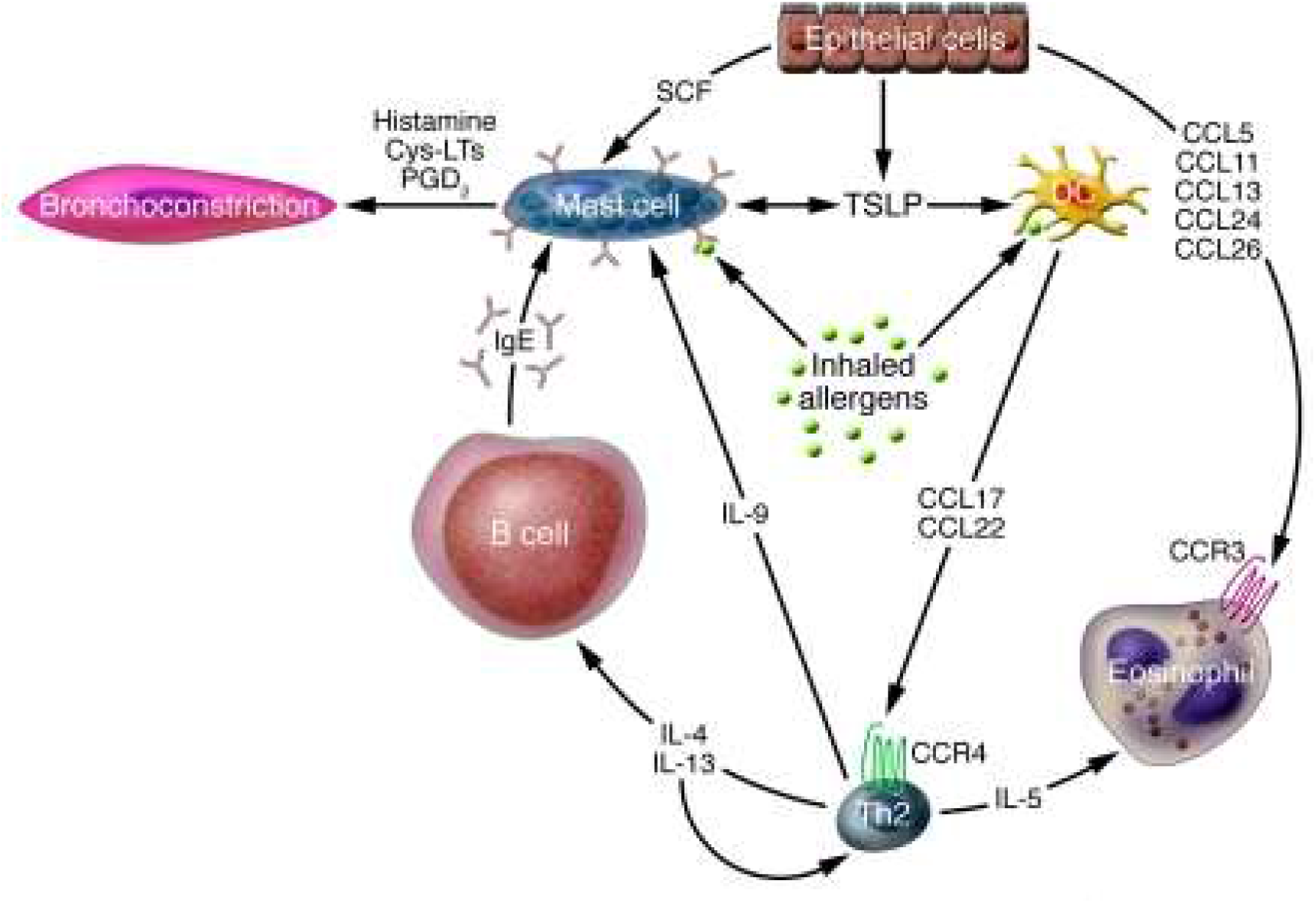
Depicts the cytokines involved in Asthma (Barnes, 2008).

**Figure 1.5.**
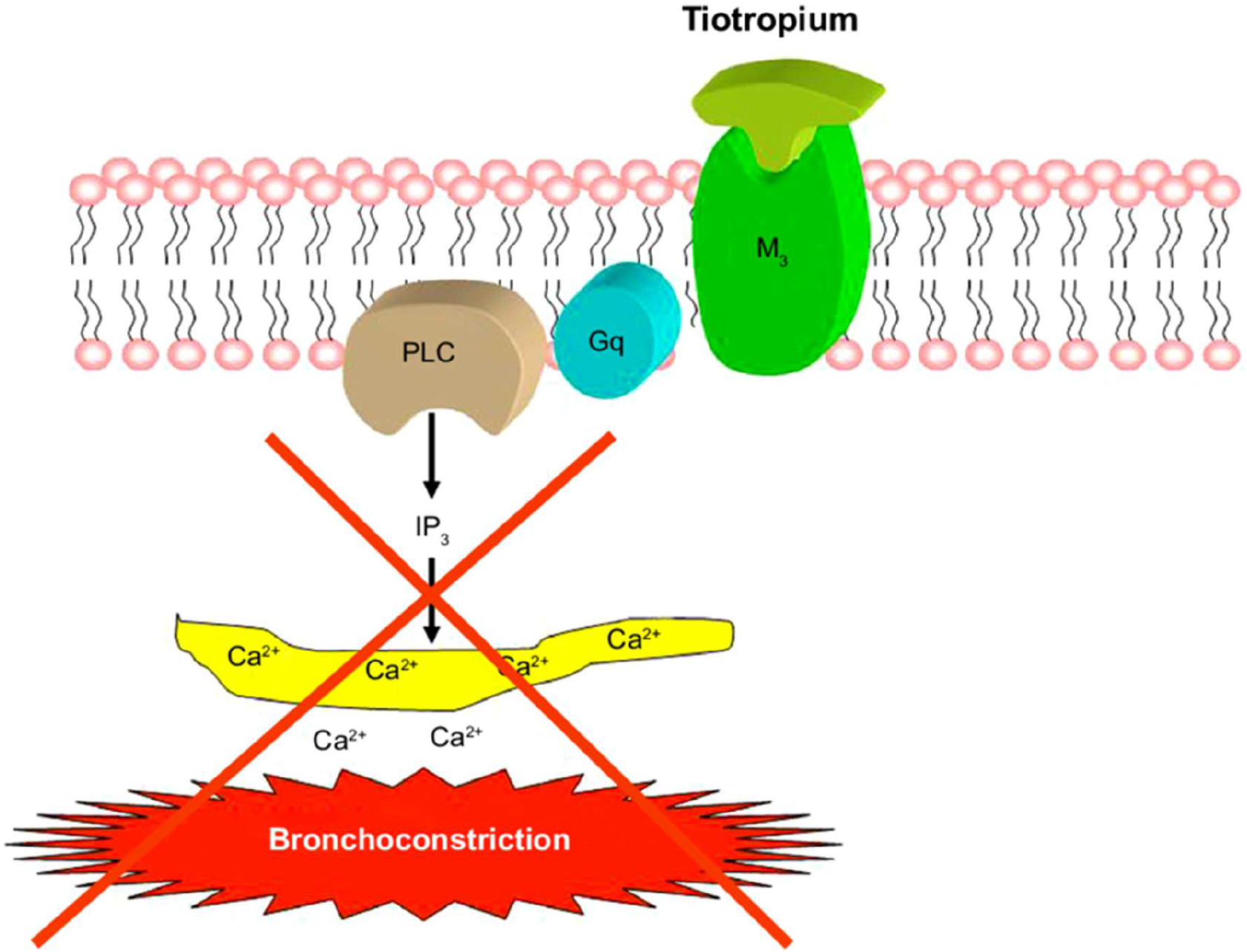
Mechanism of action of Tiotropium (Pelaia *et al.*, 2015).

Rapid dissociation from M_2_ receptor gives tiotropium a distinct kinetic selectivity. This unique selectivity is not found in SAMAs. The rapid dissociation of LAMAs generates a long-lasting effect (Lipson, 2006).

### 1.5 Human monocytic cell line THP-1 cells

The mechanistic studies of inflammatory disorders are widely carried out in *in vitro* models using human monocytes and macrophages. These monocytes clearly differentiate into macrophages at the site of inflammation (Gordon, 2007). THP-1 cells produce a number of pro-inflammatory cytokines such as IL-6, TNF- α, IL-8, IL-10. Endotoxins such as lipopolysaccharide (LPS) is frequently used to obtain inflammatory response from the cells (Gordon and Taylor, 2005). The expressions of cytokines, gene markers can be measured appropriately. The growing rate of THP-1 cells is 19-35 hours. The genetic background is responsible for its least phenotype variability which facilitates reproducibility. It is considerably high compared to commercially available peripheral blood mononuclear (PBMC)-derived cells (Chanput, Mes and Wichers, 2014). Furthermore, THP-1 monocytes are free from infections and viruses and they can be cultured for approximately 3 months and also it can be stored for years (Mangan and Wahl, 1991). Overall, THP-1 cells serve as a robust model to study the action of chemotherapeutics.

### 1.6 Enzyme Linked Immunosorbent Assay (ELISA)

Enzyme-linked immunosorbent assay (ELISA) is a plate-based immunoassay technique used to determine the levels of proteins, peptides, nucleic acids. ELISA works on the principle of antigen-antibody reactions. Enzymes such as alkaline phosphate (ALP), beta-galactosidase are commonly used (Comoglio and Celada, 1976). Antigens are immobilized on polystyrene or polyvinyl chloride plate and treated with specific antibodies. These antibodies can be identified by enzyme-labelled secondary antibodies. The antigen levels are determined by using chromogenic substrate solution. A microtiter plate reader detects the fluorescent products. ELISA is a highly specific, efficient and cost-effective assay. There are various types of ELISA such a direct, indirect, competitive, indirect competitive and sandwich ELISA (Sakamoto *et al.*, 2017). In present experiment Sandwich ELISA was performed.

### 1.7 Aims and Hypothesis

The ***specific aims*** of this project were:

1. To investigate the LPS stimulated pro-inflammatory cytokines IL-6 and TNF-α levels in THP-1 cells.
2. To investigate whether the drugs Ipratropium and Tiotropium are capable of decreasing LPS-induced inflammation in THP-1 cells.
3. To compare the effects of Ipratropium and Tiotropium in inhibiting the pro-inflammatory cytokines.

We ***hypothesize*** that Ipratropium and Tiotropium will exhibit potent anti-inflammatory effects and Tiotropium will exhibit greater anti-inflammatory action.

The alternate hypothesis was that both the therapeutic compounds would not exhibit any significant effect on the levels of IL-6 and TNF- α cytokines.

## CHAPTER 2 MATERIALS AND METHODS

Ethical approval to conduct this research experiment was obtained from Coventry University’s Ethics Committee. Current project was in complete compliance with the health and safety regulations of Coventry University. The reference number given to this project was P93689.

### 2.1 Materials

Human monocytic cell line THP-1 were acquired from the European Collection of Cell Cultures (ECACC, Health Protection Agency, Porton Down, U.K) and cultured with Rosewell Park Memorial Institute (RPMI) 1640 with Glutamax I (Invitrogen Ltd, Paisley, U.K) supplemented with 1% Penicillin Streptomycin(Pen Strep) antimycotic solution (Sigma-Aldrich) and 10% of foetal bovine serum (FBS) from Life Technologies U.K. Lipopolysaccharide (LPS) 10μl/ml, 0.05% Tween20 and 0.4% Trypan blue were purchased from Sigma-Aldrich. The drugs Fenoterol (hydrobromide) and Budesonide were obtained from Cayman Chemicals, Cambridge, U.K. Ipratropium and Tiotropium were purchased from Tocris Biosciences, Bristol, U.K. Human IL-6 DuoSet ELISA and Human TNF- α DuoSet ELISA kits were bought from R&D Systems Abingdon, U.K. TMB ELISA Substrate (High sensitivity) and 450nm Stop solution for TMB Substrate were purchased from Thermo-Fisher, U.K. Dulbecco’s Phosphate-buffered saline 1X PBS and ELISA wash buffer were bought from ThermoFisher Scientific, U.K. Lyophilized Bovine serum albumin powder was bought from Sigma-Aldrich.

### 2.2 Methods

#### Preparation of culture media

Culture media prepared had a final concentration of 10% FBS and 1% antimycotic solution. The culture media was prepared by adding 50 ml of FBS and 5.5ml of pen-strep antimycotic solution to a new 500ml bottle of RPMI. A water bath was used to warm the media to 37°C and it was then stored in a humidified atmosphere [95% air, 5%(v/v) CO_2_] at 37°C.

#### Cell culture

THP-1 cells were cultured in T-75 flasks containing 15 ml of culture media. The media was replaced every 48 hours. To replace the media, the contents of T-75 flasks (cells + media) were pipetted into a 50 ml conical-bottom centrifuge tube and it was then centrifuged at 800g for 5 minutes at room temperature. The supernatant was discarded using a graduated pipette without disturbing the cell pellet. 1ml of media was used to resuspend the cells. A cell count was performed using 10μl of cell suspension. After determining the cell count, the cell suspension was diluted to a density of 1×10^6^ cells/ml using culture media.

#### Cell counting

10μl of 0.4% trypan blue was added to 10μl of cell suspension. Using an *Improved Neubauer* Haemocytometer and a microscope, cell count was performed. The cells present in each 4×4 square of haemocytometer were counted. The volume each 4×4 square was 10-4 or 0.1μl. The dilution factor was 2. The formula for performing cell count was, cells**/**ml = cells counted in 4×4 squares x Dilution factor x 10^4^. When the cell density exceeded 1×10^6^ cells**/**ml cells were prepared for experimental use.

#### Passaging of cells

THP-1 cells double every 48 hours. Depending on the cell count, the cell suspensions were diluted appropriately to culture 15ml batches of 5 × 10^5^ cells/ml. Cell suspensions were passaged using fresh media such that the T-75 flasks have a density of 1 x 10^6^ cells/ml. To ensure that adequate number of cells were available for performing the experiment.

#### Seeding wells

On experimental days, the cells were centrifuged at 800g for 5 minutes at room temperature. The supernatant was discarded, and the pellet was resuspended in 1ml of fresh media and cell count was performed. Each well was seeded with 450μl of media containing 1 x 10^6^ cells/ml.

#### Preparation of drug solutions

Serial dilutions of stock drugs fenoterol, budesonide, ipratropium and tiotropium were performed using vehicle 1X PBS. The drugs concentrations of 10^−6^, 10^−7^ and 10^−8^ M were prepared. Requisite amount of drug solutions were prepared such that the drugs solutions would be sufficient to perform experiments in duplicates and triplicates.

#### Challenging cells with LPS

LPS vial was diluted with 1ml of PBS to prepare a concentration of 10μl/ml LPS. 25μl of LPS from reconstituted 1ml vial was added to 2475μl of PBS to make 10μl/ml concentration per 1×10^6^ cells/ml. 50μl of this concentration was used for challenging the cells. The cells were then incubated for 24 hours at 37°C (95% O_2_, 5% CO_2_ atmosphere).

#### Preparation of phosphate-buffered saline (PBS) 1X PBS

The protocol published by Wood E.J (1983) was followed and 1X PBS was prepared by transferring 8 gm of sodium chloride (NaCl), 0.2 gm of potassium chloride (KCl), 1.44 gm of disodium hydrogen phosphate Na2HPO4 and 0.24gm of potassium dihydrogen phosphate (KH_2_PO_4_) into 1000ml of distilled water. The final concentration of PBS was 137mM of NaCl, 2.7mM of KCl, 10mM of Na_2_HPO_4_ and 1.8mM of KH_2_PO_4_. The pH was adjusted to 7.2 to 7.4 using hydrochloric acid and sodium hydroxide as necessary. The solution was then sterilized by autoclaving at 121°C for 30 minutes. Later PBS was stored at room temperature. (Wood, 1983)

#### Preparation of reagent diluent (1% BSA in PBS)

5 grams of BSA was dissolved in a 500ml of 1X PBS. A magnetic stirrer was used to completely dissolve BSA. The reagent diluent (1% BSA in PBS) was stored at 4°C.

#### Preparation of ELISA wash buffer

ELISA wash buffer (0.05% Tween 20 in PBS) was prepared by adding 500μl of Tween 20 to 1000ml of sterile 1X PBS and it was mixed thoroughly. The pH was adjusted to 7.2-7.4.

#### 2.2.1 REAGENT PREPARATION

The reagents of Human IL-6 DuoSet ELISA and Human TNF-α DuoSet ELISA kits were taken and working dilutions were prepared. All the reagents were brought to room temperature before use. The reagents of each kit were sufficient to perform 5 plate assays. One-fifth of each reconstituted vial was used for a single 96 well plate assay as in accordance with the manufacturer’s instructions (R&D Systems,2019). All the following steps for ELISA assays were in accordance with the manufacturer’s instructions (R&D Systems,2019).

##### IL-6 ELISA Kit reagents

Streptavidin conjugated horseradish-peroxidase (Streptavidin-HRP): 400μl from the 2ml vial of streptavidin-HRP was added to 9600μl of reagent diluent.

##### Mouse anti-Human IL-6 capture antibody

The vial was reconstituted with 0.5ml PBS. 100μl of this reagent was added to 9600μl of reagent diluent.

##### Biotinylated goat anti-human IL-6 detection antibody

The vial was reconstituted with 1000μl of reagent diluent. 200μl of the reconstituted detection antibody was diluted to working concentration using 9600μl of reagent diluent.

##### Recombinant human IL-6 standard

The vial was reconstituted with 0.5ml of deionized water. A seven-point standard 2-fold dilutions were prepared. The stock concentration of reconstituted vial was 600 pg/ml. 500μl of reagent diluent was taken in seven serially labelled Eppendorf’s. 100μl of reconstituted standard IL-6 was transferred to first point Eppendorf and mixed thoroughly. 500μl from the first point Eppendorf was then serially transferred and mixed into remaining Eppendorf’s and so on. Overall, 100μl of standard IL-6 was taken in their respective concentrations and they were added to the 96 well enzyme immunosorbent assay (EIA) plate.

##### TNF- α ELISA kit reagents

Streptavidin conjugated horseradish-peroxidase (Streptavidin-HRP): 400μl from the 2ml vial of streptavidin-HRP was added to 9600μl of reagent diluent.

##### Mouse anti-Human TNF- α capture antibody

The vial was reconstituted with 0.5ml PBS. 100μl of this reagent was added to 9600μl of reagent diluent.

##### Biotinylated goat anti-human TNF- α detection antibody

The vial was reconstituted with 1000μl of reagent diluent. 200μl of the reconstituted detection antibody was diluted to working concentration using 9600μl of reagent diluent.

##### Recombinant human TNF- α standard

The vial was reconstituted with 0.5ml of deionized water. A seven-point standard 2-fold dilutions were prepared. The stock concentration of reconstituted vial was 10,000 pg/ml. 500μl of reagent diluent was taken in seven serially labelled Eppendorf’s. 100μl of standard stock solution was transferred to the first point Eppendorf and mixed thoroughly. 500μl from this Eppendorf was then serially transferred and mixed into the remaining Eppendorf’s and so on. Overall 100μl of standard TNF-α in their respective concentrations were added to 96 well EIA.

#### 2.2.2 PREPARATION OF ELISA IMMUNO ASSAY PLATES (EIA) PLATES

The 96 well EIA plates for IL-6 and TNF- α assay were prepared using the reagents provided in each separate kit. The reagents were separate, but the plating techniques was exactly same. The working concentration and dilutions are mentioned in the reagent preparation section.

##### Coating plates with capture antibody

The plates were coated by transferring 100μl of diluted capture antibody into 96 well plates. It was then covered with adhesive strip and stored in a refrigerator at 4°C for 12 hours.

##### Blocking plates

After coating the plates with capture antibody, the plates were washed using ELISA wash buffer (0.05% Tween20 in PBS). Firstly, the adhesive strip was removed, and contents of the plate were discarded. The adhesive strip was retained. Each well was filled with wash buffer and the plate was allowed to rest for 30 seconds. The wash buffer was discarded, and the wells were filled with wash buffer again. The washing step was repeated twice. After discarding wash buffer, the plates were inverted and gently tapped on absorbent paper to ensure that all the wells were dry and free from air bubbles.300μl of reagent diluent was added to each well. The plate was then covered with the retained adhesive strip and allowed to incubate at room temperature for 2 hours.

##### Addition of standards and samples

Respective standards IL-6, TNF- α and the cell culture supernatant samples were added to the enzyme immunoassay plate (EIA plate) and incubated for 2 hours at room temperature.

##### Incubation with detection antibody

100μl of detection antibody diluted previously in reagent diluent was transferred in each well and covered with adhesive strip. The plate was allowed to incubate for 2 hours at room temperature.

##### Incubation with Streptavidin-HRP

100μl of Streptavidin-HRP diluted in reagent diluent was transferred into each well of the plate. It was covered with adhesive strip and incubated for 20 minutes in a dark place.

##### Addition of TMB ELISA Substrate solution

100μl substrate solution was added to each well and the plate was incubated for 20 minutes at room temperature in a dark place.

##### Addition of TMB ELISA Stop solution

50μl of stop solution was added to each well and the plate was tapped gently to ensure complete mixing of the solution.

#### 2.2.3 MEASUREMENT OF OPTICAL DENSITY (ABSORBANCE) OF EIA PLATES

Labtech LT-4500 microplate reader (Labtech, 2019) was used to measure absorbance of each well. The wavelength of the instrument was set at 450nm. Wavelength correction was set to 450nm - 540nm. A standard curve was generated for the standards and also the test drugs to determine the cytokine concentrations.

### 2.3 RESEARCH PROTOCOL

Evaluation of LPS induced cytokine production in THP-1 cell culture supernatants by using ELISA

THP-1 cells were taken in 24 well plate. Each well was seeded with 450μl of media containing 1 × 10^6^ cells/ml. These cells were challenged with 10μl/ml LPS and incubated for 12 hours. The cells were then treated with fenoterol, budesonide, ipratropium and tiotropium in concentrations of 10^−6^,10^−7^ and 10^−8^ M for 60 minutes. The cell suspensions (LPS + cells + Drugs) were transferred in labelled centrifuge tubes and then they were centrifuged for 5 minutes at 800g. The supernatants were collected and used for measurement of IL-6 and TNF- α levels using separate DuoSet ELISA kits (R&D Systems).

For performing ELISA, the plates were first coated with 100μl capture antibodies and then incubated overnight at room temperature. The plates were washed using ELISA wash buffer (0.05% Tween20 in PBS) and then the plates were blocked using 300μl of reagent diluent (1% BSA in PBS) and the plate was incubated at room temperature for 2 hours. The plates were then washed using the wash buffer and 100μl of culture supernatants and standard reagents IL-6 and TNF- α were added and incubated for 2 hours at room temperature. After incubation the plates were washed and 100μl of streptavidin-HRP was added. The plates were incubated for 20 minutes in a dark place at room temperature. The plates were washed again and 100μl of TMB ELISA substrate solution was added to each to each well and allowed to incubate for another 20 minutes in a dark place. 50μl of TMB ELISA stop solution was transferred to the plates. The optical density of each plate IL-6 and TNF- α was measured using microplate reader set at 450nm wavelength. The cytokine concentrations (pg/ml) obtained were calculated and compared with the standards IL-6 and TNF- α.

### 2.4 STATISTICAL ANALYSIS

Experimental data was presented as mean ± S.E.M. Statistical analysis was performed for “N” number of observations. Students *t-*test, Mann-Whitney *U* test and one-way ANOVA statistic tests were used for statistical analysis. Bonferroni’s *post-hoc* test was performed to compare 3 or more data groups. The statistical significance was considered only when *P*<0.05 between the groups. GraphPad Prism 8 computer software (Graphpad.com, 2018) was used to analyse and interpret the experimental data.

## CHAPTER 3 RESULTS

### 3.1 ELISA results for IL-6

The production of pro-inflammatory cytokine IL-6 was assessed using ELISA. THP-1 cells were treated with LPS to determine the IL-6 cytokine concentration and they were compared with untreated cells. The experimental data was expressed as mean ± S.E.M. The ELISA data for IL-6 concentration (pg/ml) was confirmed by constructing a standard curve using IL-6 standards provided in DuoSet IL-6 ELISA kit (data not shown). There was no significant difference between LPS negative control and the vehicle control (PBS) (data not shown).

#### 3.1.1 Comparison of IL-6 production in LPS treated and control (untreated) THP-1 cells

As shown in figure 3.1, a substantial increase in the IL-6 cytokine concentration was observed in LPS treated cells. The mean ± SEM data obtained was 262.85 ± 1.7 pg/ml for LPS treated cells and 15.4 ± 0.8 for control. The control (untreated) cells showed least concentration 15 pg/ml whereas the LPS treated cells had 262.85 pg/ml of IL-6. The data represented in the figure confirms that IL-6 levels increased significantly in LPS treated cells. The experimental technique was thus validated.

**Figure 3.1.**
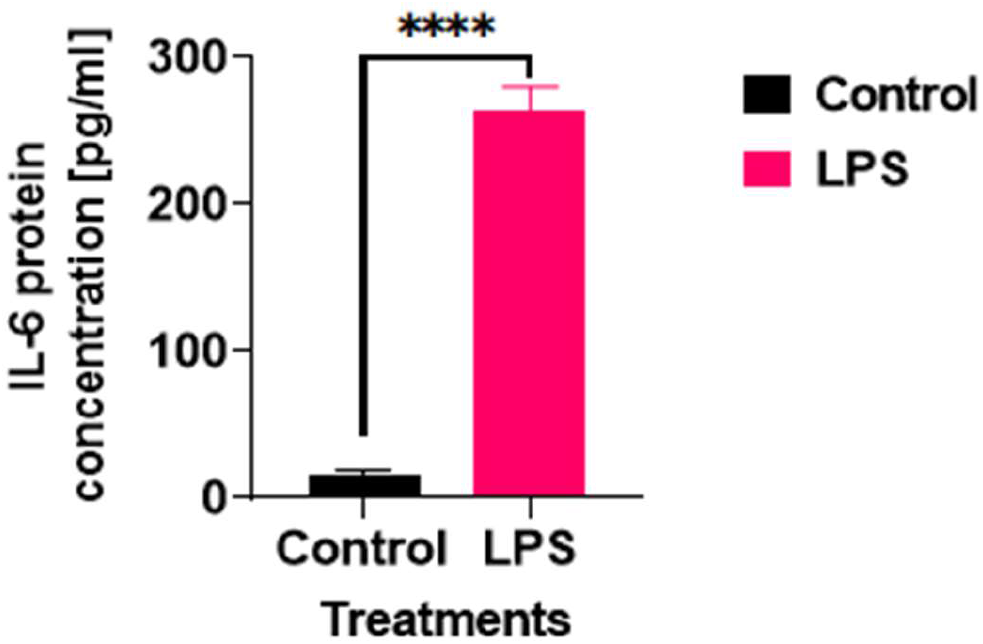
A comparison of control (untreated) and LPS treated cells THP-1 cells. The data was represented as mean ± S.E.M for N= 4 for both groups and the experiment was performed in duplicates. **** = P<0.0001 vs. control and it indicates that a statistically significant difference is present between the treated and untreated cells.

*Note - All the figure illustrated in this report are Colour-blind Safe.*

#### 3.1.2 Determination of the effect of Budesonide on IL-6 protein expression in LPS stimulated THP-1 cells

Budesonide and Fenoterol were used a positive control. Budesonide is an existing anti-inflammatory drug which is known for its cytokine reducing properties. As outlined in 3.1 IL-6 data was validated by generating a standard curve. The production of IL-6 cytokine in the cells was stimulated by using LPS and the cells treated with drugs were compared with untreated (LPS only) cells. The concentrations of Budesonide used were 1 × 10^−6^ M, 1 × 10^−7^ M and 1 × 10^−8^ M. The mean ± S.E.M data obtained for untreated control (LPS only) cells was 262.85 + 1.7 pg/ml. For the cells treated with various concentrations of Budesonide such as 1 × 10^−8^ M, 1 × 10^−7^ M and 1 × 10^−6^ M, the mean ± SEM data was 149.31 ± 3, 100.4 ± 2.5 and 90.01 ± 2.2 pg/ml.

As shown in figure 3.2, the concentration 1 × 10^−6^ M was found to be nearly as potent as 1 × 10^−7^ M. The IL-6 concentrations (pg/ml) gradually decreased in the cells treated with Budesonide in a statistically significant dose-responsive manner.

**Figure 3.2.**
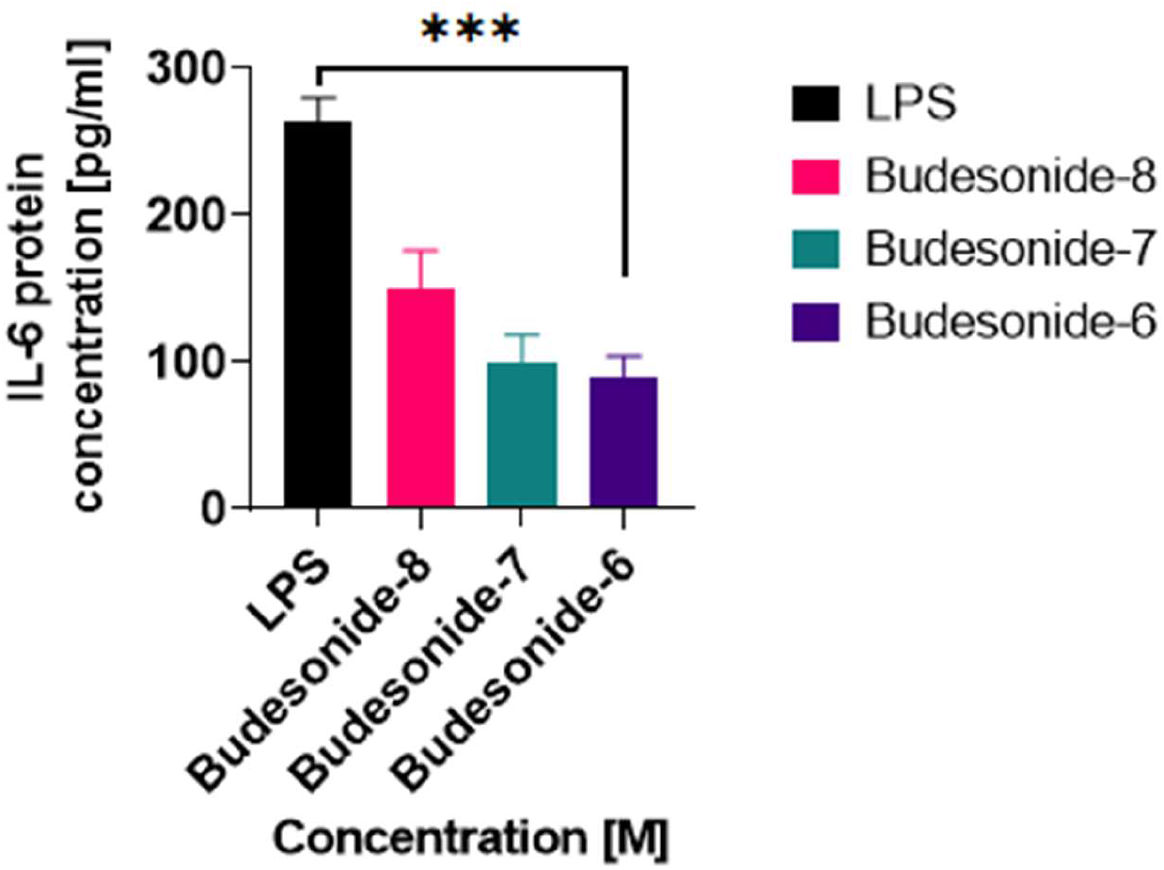
Graphical representation of the effect of Budesonide on IL-6 concentration in LPS stimulated THP-1 cells. The data was presented as mean ± S.E.M for N= 4 for all the groups. ***= P<0.001 vs. LPS control indicates that a statistically significant difference is present.

#### 3.1.3 Determination of the effect of Fenoterol on IL-6 protein expression in LPS stimulated THP-1 cells

Fenoterol is an existing anti-inflammatory drug which is known for its cytokine reducing properties. It was used as a positive control. As outlined in 3.1 IL-6 data was validated by generating a standard curve. The concentrations of Fenoterol used were 1 × 10^−6^ M, 1 × 10^−7^ M and 1 × 10^−8^ M. The mean ± S.E.M data obtained for untreated control (LPS only) cells was 262.85 + 1.7. For the cells treated with various concentrations of Fenoterol 1 × 10^−8^ M, 1 × 10^−7^ M and 1 × 10^−6^ M, the mean ± SEM data obtained was 233.53 ± 6.2, 154.5 ± 5.0 and 111.8 ± 3.8 pg/ml.

The *P* value was found to be *P<0.001* vs. LPS control for all the groups, indicating that there was a statistical difference present between the groups. As shown in figure 8 the IL-6 concentrations in LPS stimulated THP-1 cells gradually declined after treatment with Fenoterol.

#### 3.1.4 Determination of the effect of Ipratropium on IL-6 protein expression in LPS stimulated THP-1 cells

Ipratropium was used as a test drug to assess its IL-6 cytokine reducing properties in present *in vitro* study. The concentrations used were 1 × 10^−6^ M, 1 × 10^−7^ M and 1 × 10^−8^ M. The mean ± S.E.M data obtained for untreated control (LPS only) cells was 262.85 + 1.7. For the cells treated with various concentrations of Ipratropium such as 1 × 10^−8^ M, 1 × 10^−7^ M and 1 × 10^−6^M, the mean ± SEM data obtained was 233.91 ± 3.62, 236.26 ± 2.9 and 166.9 ± 3.3 pg/ml.

As shown in figure 3.4, there was a decline in IL-6 concentration in Ipratropium treated cells compared to untreated LPS only cells, but statistically the results were not significant. *P* value for all data sets was found to be *P>0.01* vs LPS control.

**Figure 3.3.**
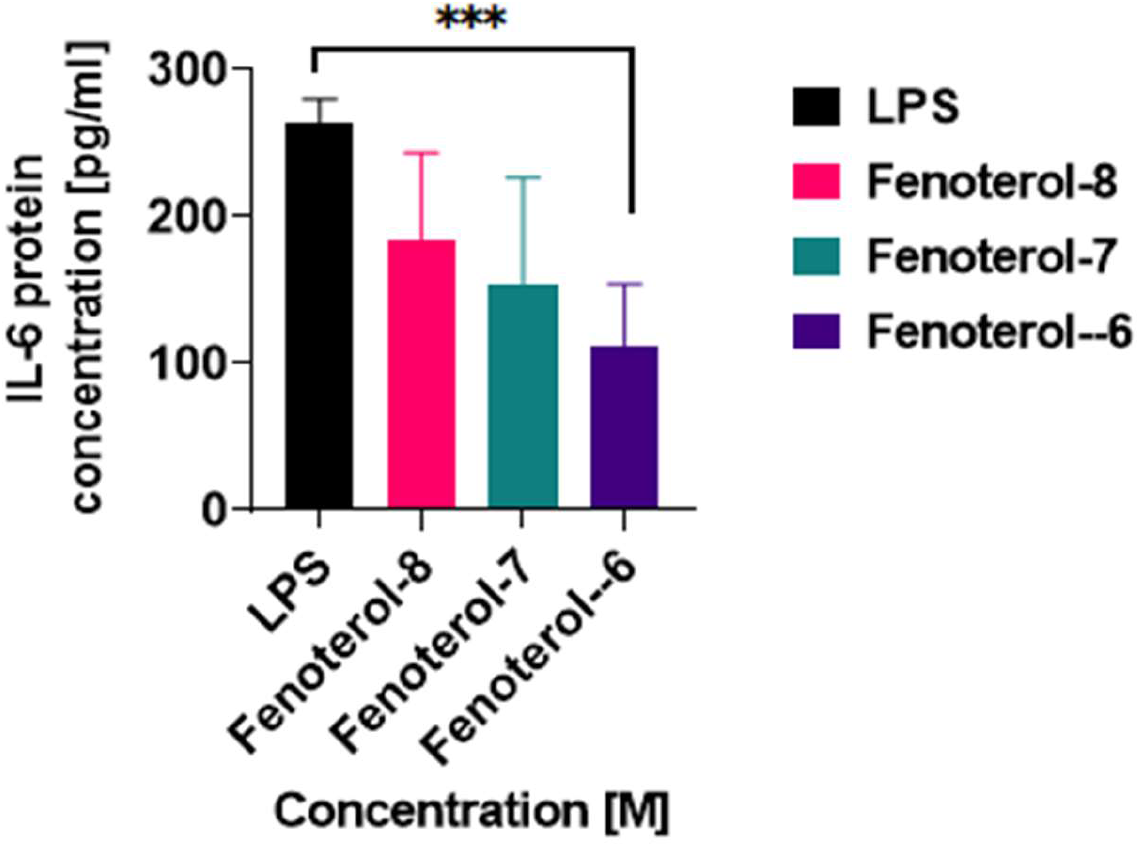
The effect of Fenoterol on IL-6 protein concentration in LPS stimulated THP-1 cells. The data was presented as mean ± S.E.M for N = 4 for all the groups. ***= P<0.001 vs. LPS control indicates that a statistically significant difference is present.

**Figure 3.4.**
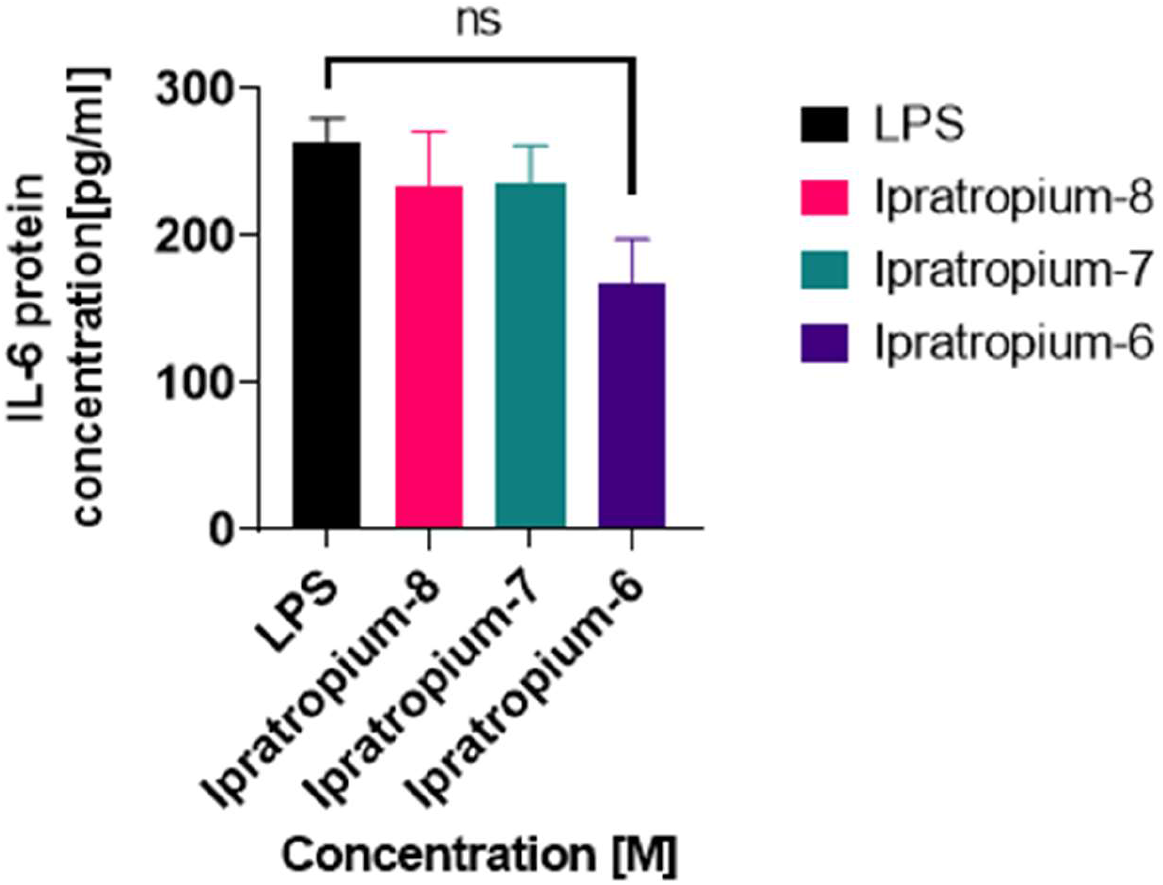
The effect of Ipratropium on IL-6 protein concentration in LPS stimulated THP-1 cells. The data was presented as mean ± S.E.M for N = 4 for all the groups. ns = P>0.01 vs. LPS control, indicates that there was no statistically significant difference in the data groups.

#### 3.1.5 Determination of the effect of Tiotropium on IL-6 protein expression in LPS stimulated THP-1 cells

Tiotropium was used as a test drug to assess its IL-6 reducing properties in present *in vitro* study. The concentrations used were 1 × 10^−6^ M, 1 × 10^−7^ M and 1 × 10^−8^ M. The mean ± S.E.M data obtained for untreated control (LPS only) cells was 262.85 ± 1.7 pg/ml. For the cells treated with various concentrations of Tiotropium such as 1 × 10^−8^ M, 1 × 10^−7^ M and 1 × 10^−6^ M, the mean ± SEM data obtained was 243.77 ± 3.6, 158.57 ± 4.03 and 156.3 ± 3.2 pg/ml.

Figure 3.5 graphically represents the moderate decrease in the IL-6 concentrations after treatment with Tiotropium. Both 1 × 10^−7^ M, 1 × 10^−6^ M Tiotropium concentrations effectively reduced nearly 50% of the IL-6 concentration. *P* value obtained for all the data bars was found to be *P*<0.05 vs LPS control.

**Figure 3.5.**
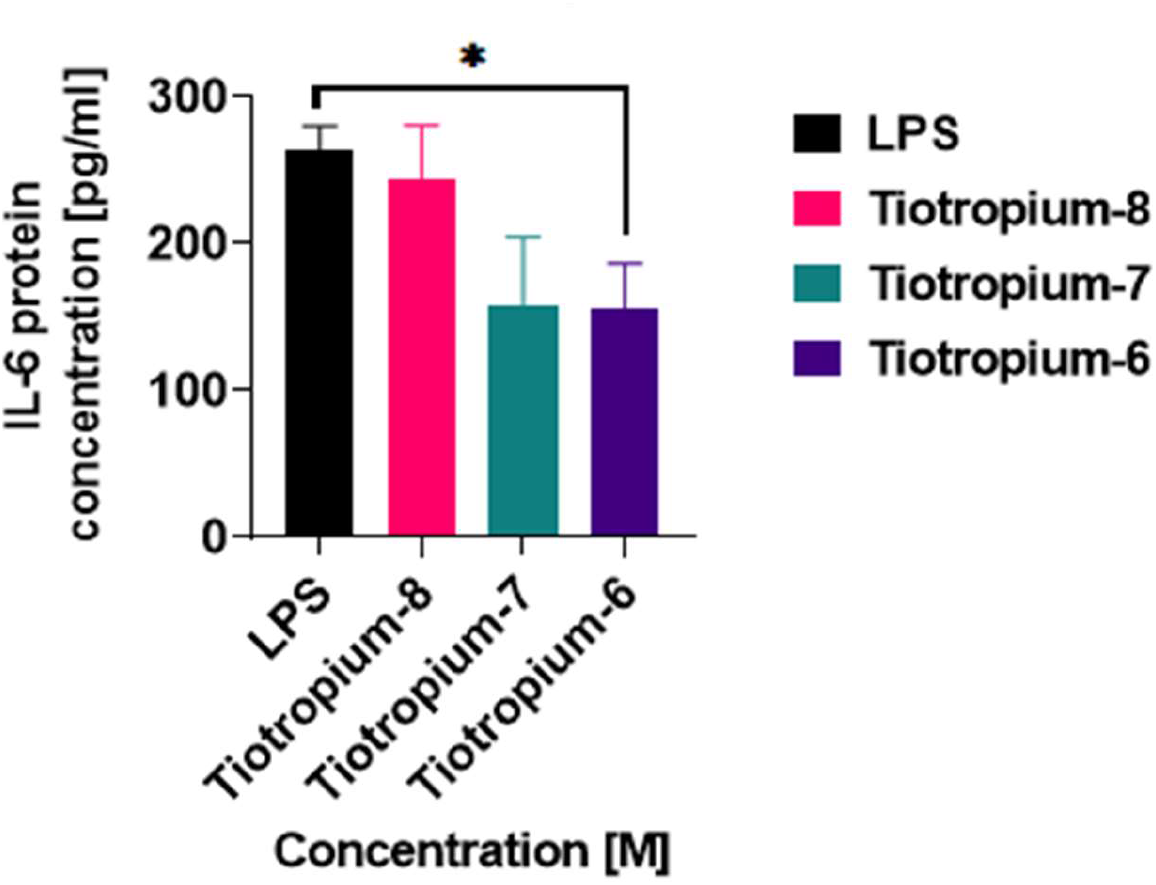
The effect of Tiotropium on IL-6 protein concentration in LPS stimulated THP-1 cells. The data was presented as mean ± S.E.M for N = 4 for all the groups. *\** = P<0.05 vs. LPS control, it indicates that there is a statistical difference between the groups.

### 3.2 ELISA results for TNF- α

The production of pro-inflammatory cytokine TNF- α was assessed using ELISA. THP-1 cells were treated with LPS to determine the TNF- α cytokine concentration and they were compared with untreated cells. The experimental data was expressed as mean ± S.E.M. The ELISA data for TNF- α concentration (pg/ml) was confirmed by constructing a standard curve using TNF- α standards provided in DuoSet TNF- α ELISA kit (data not shown). There was no significant difference between LPS negative control and the vehicle control (PBS) (data not shown).

#### 3.2.1 Comparison of TNF- α production in LPS treated and control (untreated) THP-1 cells

As shown in figure 3.6, a substantial increase in the TNF- α cytokine concentration was observed in LPS treated cells. The mean ± SEM data obtained was 726.33 ± 3.7 pg/ml for LPS treated cells and 42.03 ± 0.8 pg/ml for control (untreated cells). The control showed least concentration 42 pg/ml of TNF- α, whereas the LPS treated cells had a concentration of 726 pg/ml. The data represented in figure 11 confirms that TNF- α levels increased significantly in LPS treated cells. The research methodology was thus validated.

**Figure 3.6.**
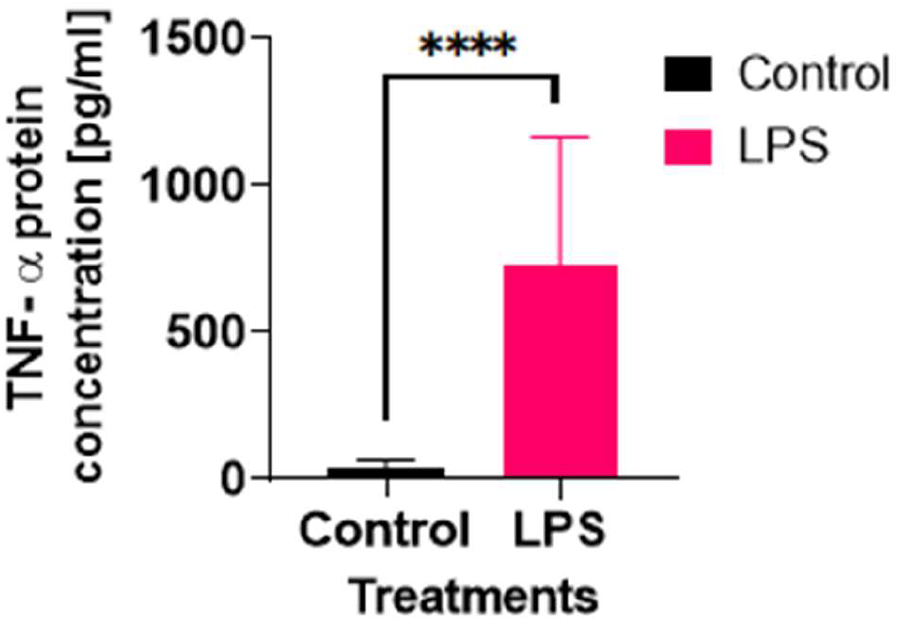
A comparison of control (untreated) and LPS treated cells THP-1 cells. The data was represented as mean ± S.E.M for N= 4 for both groups and the experiment was performed in duplicates. **** = P<0.0001 vs. control and it indicates that a statistically significant difference is present between the treated and untreated cells.

#### 3.2.2 Determination of the effect of Budesonide on TNF- α protein expression in LPS stimulated THP-1 cells

Budesonide and Fenoterol were used a positive control. Budesonide is an existing anti-inflammatory drug which is known for its cytokine reducing properties. As outlined in 3.2 TNF- α data was validated by plotting a standard curve. The production of TNF- α cytokine in the cells was stimulated by using LPS and the cells treated with drugs were compared with untreated (LPS only) cells. The concentrations of Budesonide used were 1 × 10^−6^ M, 1 × 10^−7^ M and 1 × 10^−8^ M. The mean ± S.E.M data obtained for untreated control (LPS only) cells was 736.33 ± 3.7 pg/ml. For the cells treated with various concentrations of budesonide such as 1 × 10^−8^ M, 1 × 10^−7^ M and 1 × 10^−6^ M, the ± SEM data was 568.60 ± 8.2, 252.92 ± 5.1 and 144.94 ± 3.6 pg/ml.

As depicted in figure 3.7 Budesonide decreased the TNF- α levels considerably. A three-fold reduction in TNF- α concentration was observed in the LPS stimulated cells treated with budesonide concentration 1 × 10^−6^ M.

**Figure 3.7.**
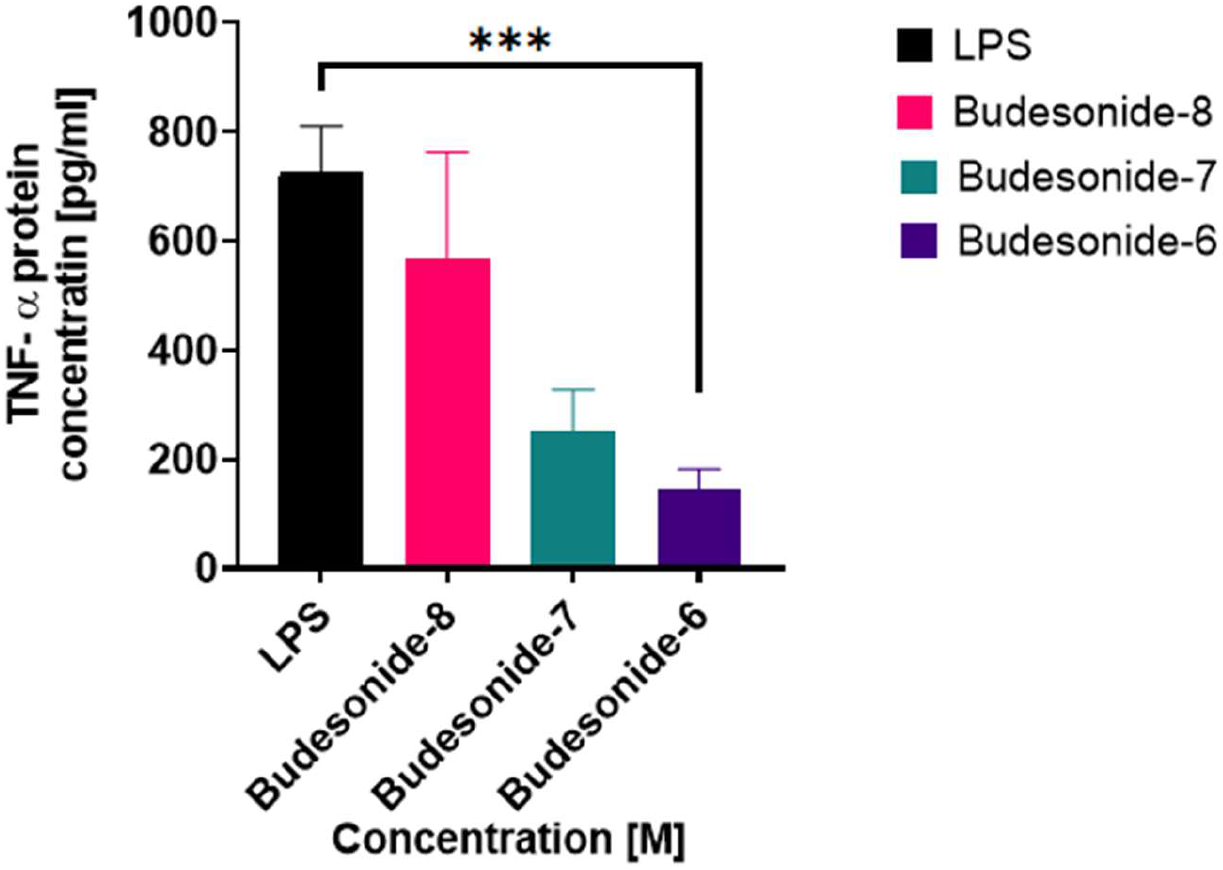
Graphical representation of the effect of Budesonide on TNF- α concentration in LPS stimulated THP-1 cells. The data was presented as mean ± S.E.M for N= 4 for all the groups. ***= P<0.001 vs. LPS control, indicates that a statistically significant difference is present between the groups.

#### 3.2.3 Determination of the effect of Fenoterol on TNF- α protein expression in LPS stimulated THP-1 cells

Fenoterol was used as a positive control. The mean ± SEM experimental data for LPS only cells (control) was 736.33 ± 3.7 pg/ml and for various concentrations of fenoterol such as 1 × 10^−8^ M, 1 × 10^−7^ M and 1 × 10^−6^ M was it found to be 474.17 ± 6.5, 292.42 ± 5.8 and 144.9 ± 3.6 pg/ml. The *P* value was found to *P*<0.001 vs LPS control for all the groups. It indicated that there was a statistically significant difference between the groups.

As shown in figure 3.8, there was a steady decline in TNF- α protein expression in Fenoterol treated cells. The responses were in dose-response pattern. A three-fold reduction in cytokine expression was observed in cells treated with the concentration 1 × 10^−6^ M.

**Figure 3.8.**
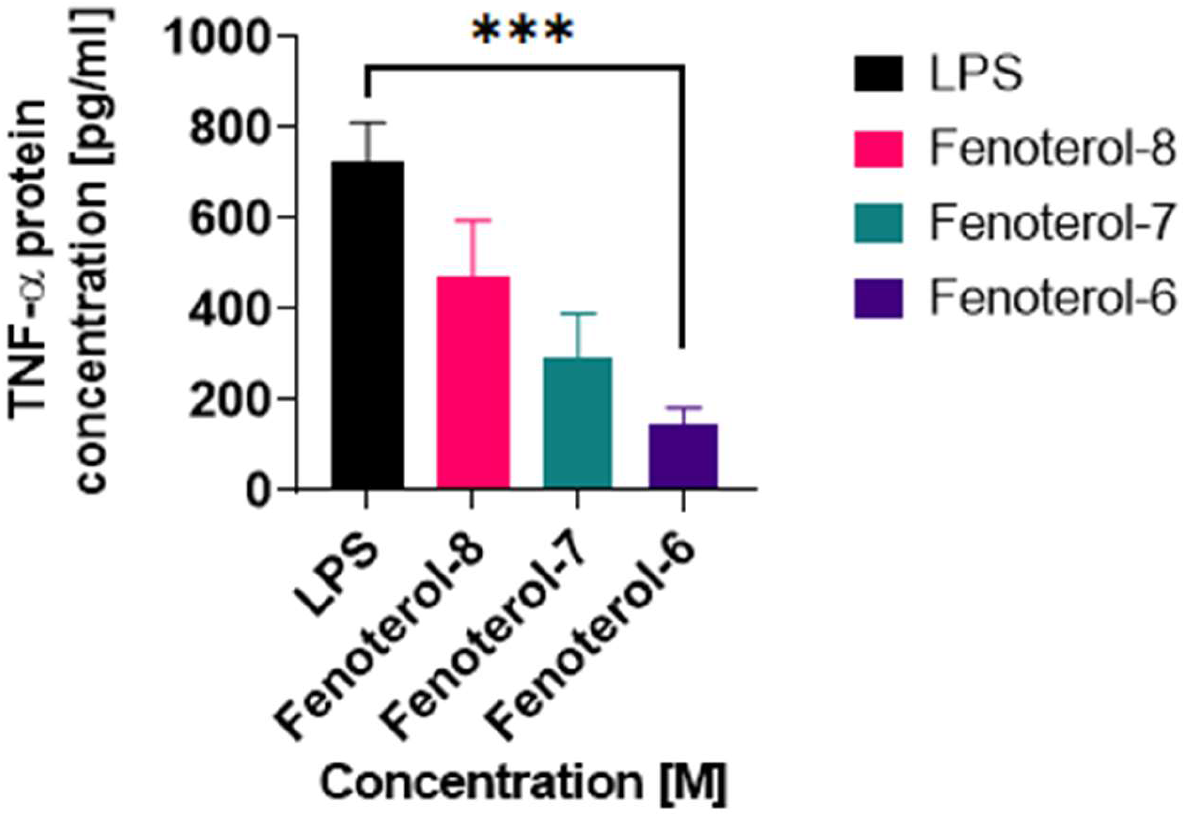
Graphical representation of the effect of Fenoterol on TNF- α concentration in LPS stimulated THP-1 cells. The data was presented as mean ± S.E.M for N =4 for all the groups. ***= P<0.001 vs. LPS control indicates that a statistically significant difference is present between the groups.

#### 3.2.4 Determination of the effect of Ipratropium on TNF- α protein concentration in LPS stimulated THP-1 cells

Ipratropium was used as a test drug to examine its cytokine TNF- α reducing ability in present *in vitro* study. The concentrations used were 1 × 10^−6^ M, 1 × 10^−7^ M and 1 × 10^−8^ M. The mean ± S.E.M data obtained for untreated control (LPS only) cells was 736.33 ± 3.7pg/ml. For the cells treated with various concentrations of Ipratropium such as 1 × 10^−8^ M, 1 × 10^−7^ M and 1 × 10^−6^ M, the mean ± SEM data obtained was 562.05 ± 58, 531.34 ± 6.3 and 400.6 ± 5.71 pg/ml. The *P* value was found to be *P*>0.01 vs. LPS control for all the groups.

There was a slight decrease in the concentration of TNF- α in cells treated with Ipratropium but mathematically these observations were non-significant. Figure 3.9 illustrates that the observations were not statistically significant.

**Figure 3.9.**
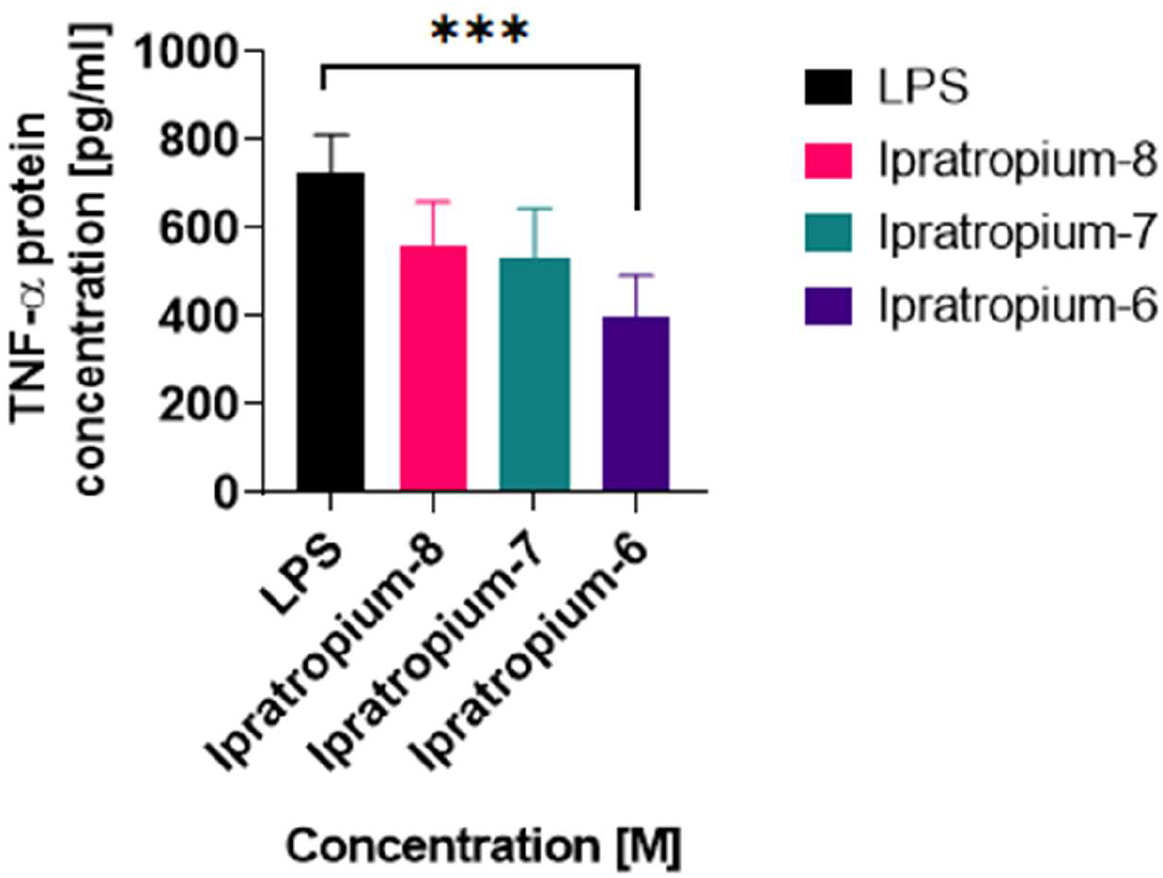
Graph depicting the effect of Ipratropium on TNF- α protein concentration in LPS stimulated THP-1 cells. The data was presented as mean ± S.E.M for N = 4 for all the groups. ns = P>0.01 vs. LPS control indicates that there was no statistically significant difference between the data groups.

#### 3.2.5 Determination of the effect of Tiotropium on TNF- α protein concentration in LPS stimulated THP-1 cells

Tiotropium was also used as a test drug to assess its TNF- α reducing properties in present *in vitro* study. The concentrations used were 1 × 10^−6^ M, 1 × 10^−7^ M and 1 × 10^−8^ M. The mean ± S.E.M data obtained for untreated control (LPS only) cells was 736.33 ± 3.7 pg/ml. For the cells treated with various concentrations of Tiotropium such as 1 × 10^−8^ M, 1 × 10^−7^ M and 1 × 10^−7^ M, the mean ± SEM data obtained was 226.54 ± 4.4, 192.61 ± 3.8 and 207.48 ± 5.08 pg/ml.

Figure 3.10 demonstrates that approximately 70% of the TNF- α protein concentration declined in the cells treated with Tiotropium. The *P* value obtained was *P*<0.0001 vs. LPS control for all the groups, which indicates that the results obtained were highly statistically significant. Overall a considerable decrease in the cytokine expression was observed in cells treated with Tiotropium.

**Figure 3.10.**
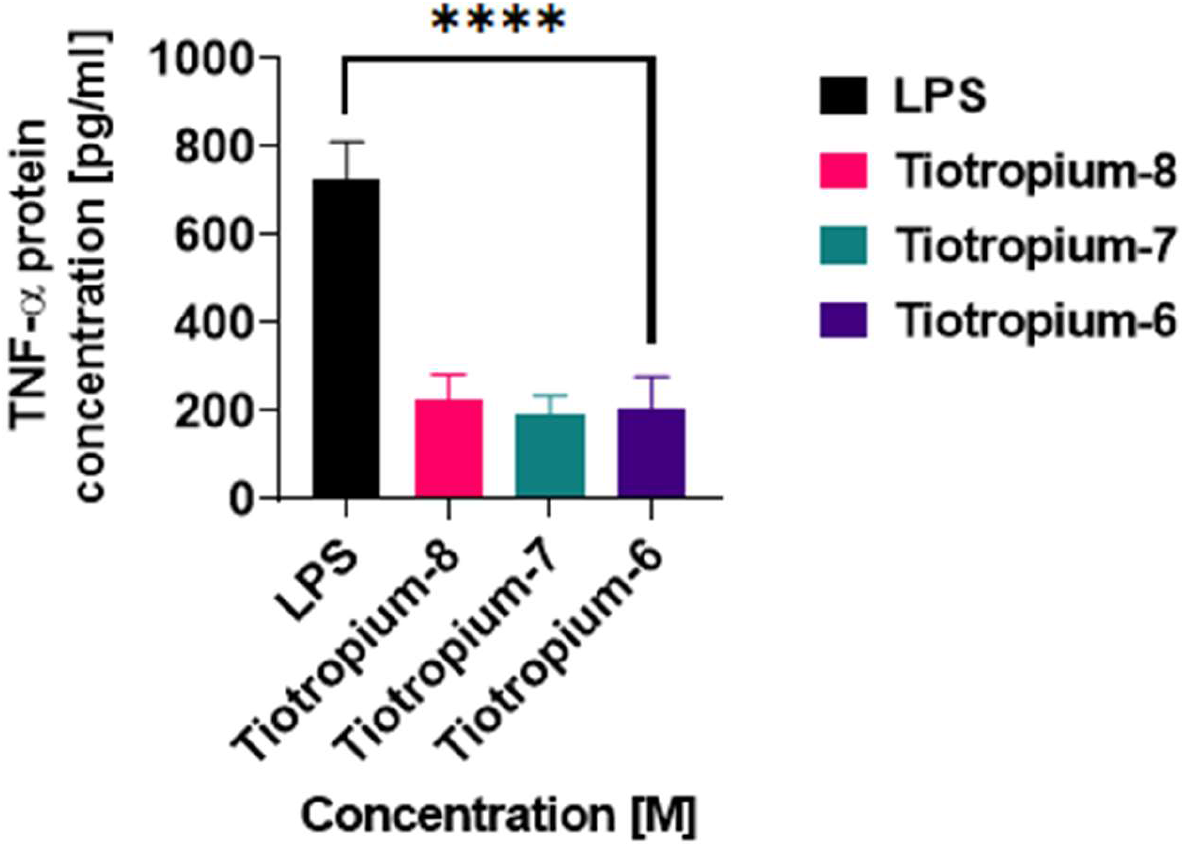
The effect of Tiotropium on TNF-α protein concentration in LPS stimulated THP-1 cells. The data was presented as mean ± S.E.M for N = 4 for all the groups. ****= P<0.0001 vs. LPS control indicates that a statistically significant difference is present between the groups.

### 3.3 Quantitative inhibition of IL-6 concentration following the drug treatments

Budesonide, Fenoterol, Ipratropium and Tiotropium of varying concentrations 1 × 10^−6^ M, 1 × 10^−7^ M and 1 × 10^−8^ M were used to treat LPS-stimulated THP-1 cells. The baseline or highest relevancy point for IL-6 was set at 262.85 pg/ml. This value represents the maximum cytokine concentration detected in THP-1 cells subsequent to LPS challenge. Figure 3.11 illustrates the dose-responsive decline in IL-6 levels, Ipratropium and Tiotropium decreased the IL-6 levels significantly with respect to their concentration. On the contrary Budesonide and Fenoterol were found to be more efficient in reducing IL-6 levels than Ipratropium and Tiotropium. *P* value was found to be *P*<0.05 vs. control IL-6 baseline and it indicates that the results obtained were statistically significant.

**Figure 3.11.**
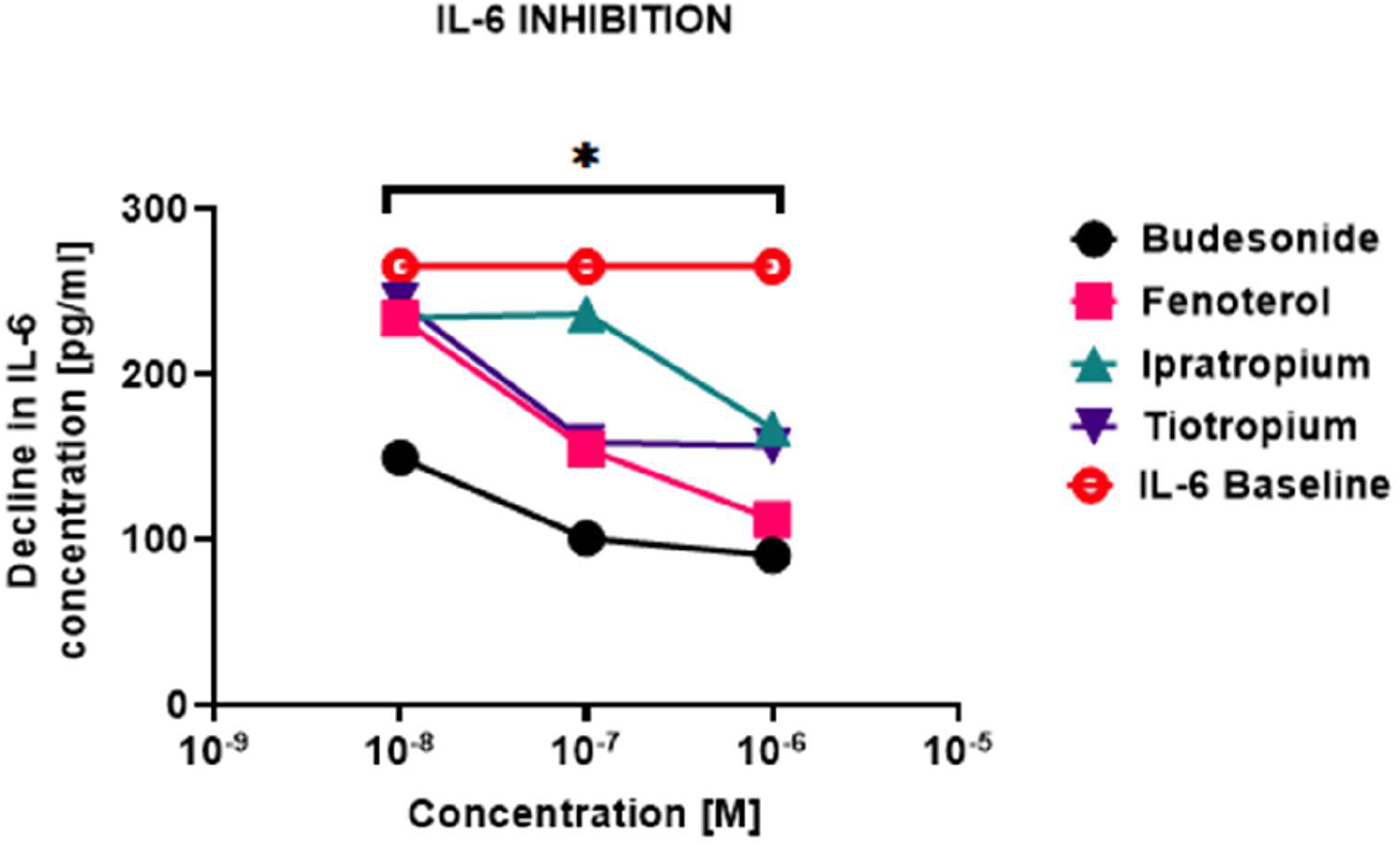
Graph depicting the effect of Budesonide, Fenoterol, Ipratropium and Tiotropium on IL-6 protein concentration in contrast with IL-6 baseline or upper control limit. The data was presented as mean ± S.E.M for N = 4 for all the groups. * = P<0.05 vs control IL-6 baseline which indicates that a statistically significant difference is present between the data groups.

Ipratropium, Tiotropium and Fenoterol at a low concentration of 1 × 10^−8^ M have minimal effect on IL-6, whereas Budesonide reduced approximately 50% of the total cytokine concentration. As shown in figure 3.11, the drugs elicited maximum effect on IL-6 expression particularly at the concentration 1 × 10^−6^ M. The results also indicate precisely, that IL-6 reduction is proportional to the concentration of the drugs.

### 3.4 Quantitative inhibition of TNF-α concentration following the drug treatments

Budesonide, Fenoterol, Ipratropium and Tiotropium of varying concentrations 1 × 10^−6^ M, 1 × 10^−7^ M and 1 × 10^−8^ M were used to treat LPS-stimulated THP-1 cells. The baseline or highest relevancy point for IL-6 was set at 736.33 pg/ml. This value represents the maximum cytokine concentration detected in THP-1 cells subsequent to LPS challenge. Figure 3.12 illustrates the dose-responsive decline in TNF- α levels. Ipratropium and Tiotropium decreased the TNF-α levels significantly with respect to their concentration. Tiotropium substantially reduced TNF-α concentration even at least concentration of 1 × 10^−6^ M, thereby it followed a linear trend. Ipratropium however reduced the cytokine levels and demonstrated a dose-responsive pattern. The *P* value was found to be *P*<0.01 vs. control TNF-α baseline and it indicates that the results obtained were statistically significant.

**Figure 3.12.**
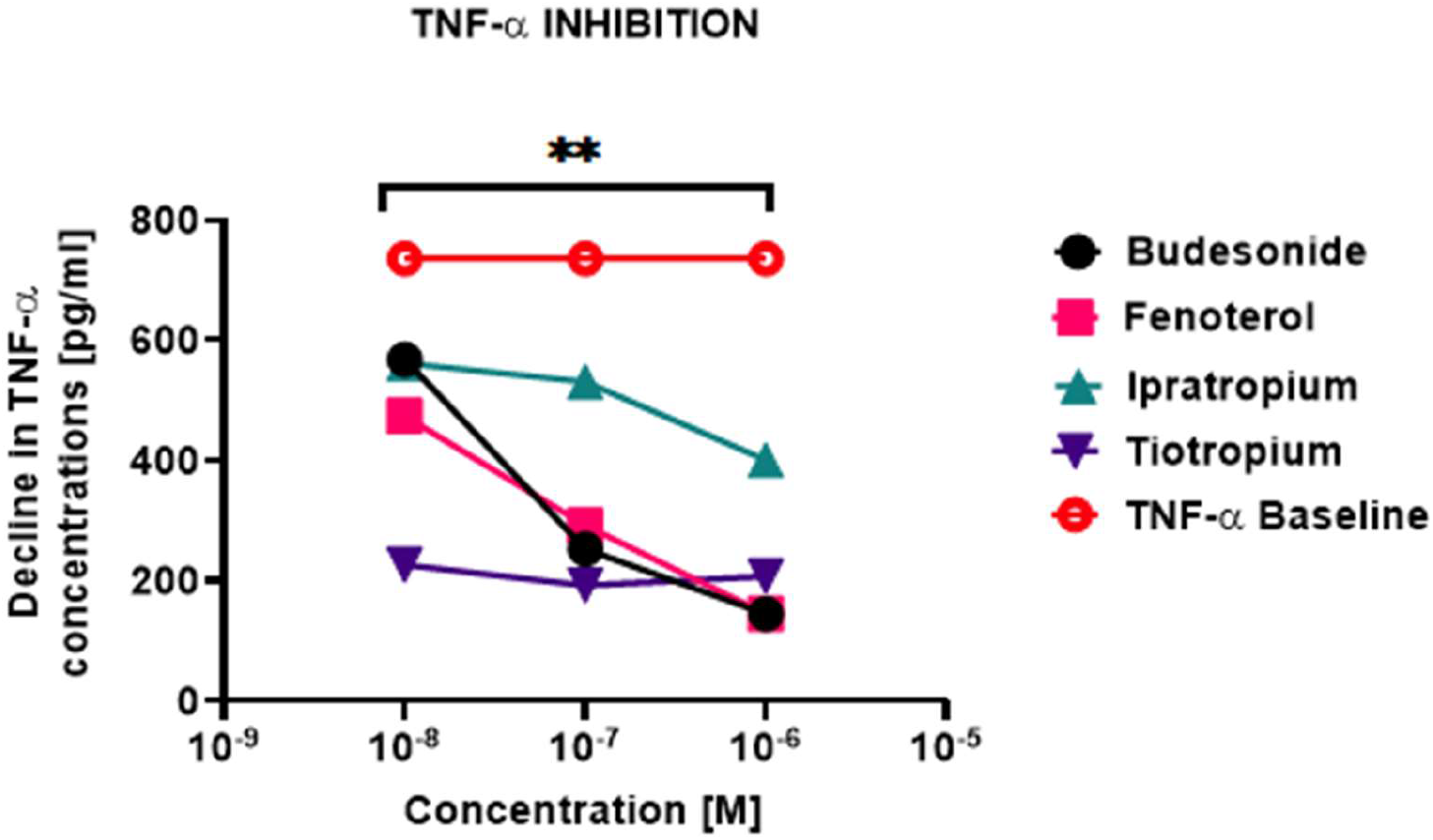
Graph depicting the effect of Budesonide, Fenoterol, Ipratropium and Tiotropium on TNF- α protein concentration in contrast with TNF- α baseline or upper control limit. The data was presented as mean ± S.E.M for N = 4 for all the groups. ** = P<0.01 vs control TNF- α baseline which indicates that a statistically significant difference is present between the data groups.

From figure 3.12 it is evident that Budesonide and Fenoterol exponentially reduced TNF- α levels in a dose-responsive manner. Additionally, they elicited identical responses and demonstrated higher TNF- α reduction potential than Ipratropium and Tiotropium. Furthermore, the results obtained were highly significant statistically.

## CHAPTER-4 DISCUSSION

### 4.1 Analysis and interpretation of the experimental findings

The experimental data confirms that Ipratropium and Tiotropium significantly decrease the expression of IL-6 and TNF- α cytokines in LPS stimulated THP-1 cells under *in vitro* conditions. The results of the present study demonstrated that the endotoxin LPS substantially elevated the cytokine expression and it was confirmed by the quantification of IL-6 and TNF-α levels using ELISA as shown in figure 3.1 and 3.2. Furthermore, the cytokine inhibiting characteristics of standard drugs Budesonide and Fenoterol were ascertained during the investigation of IL-6 and TNF- α concentrations as shown in figures 3.2, 3.3, 3.7 and 3.8. The results were statistically significant and overall the standard drugs followed a dose-responsive pattern, consequently validating the research methodology.

The inflammatory stress induced by LPS promoted TNF- α and IL-6 production, TNF-α levels were comparatively higher than IL-6 which is evident in figure 3.1 and 3.6. The reason for difference in the concentrations of cytokines is that the LPS upregulates E-selectin mRNA expression effectively which results in increased TNF- α production. LPS elevates TNF- α synergistically by activating NF-κB pathway which further amplifies the inflammatory response in cells (Jersmann *et al.*, 2001). The transcription of IL-6 mRNA is also stimulated by NF-κB signalling pathway. The activation of glucocorticoid receptors supresses the IL-6 gene expression and inhaled-corticosteroids (ICS) Budesonide follows this mechanism to reduce inflammation (Tanaka, Narazaki and Kishimoto, 2014). Figure 3.2 and 3.7 illustrate the significant decline in cytokine concentrations. Budesonide is commonly used for the treatment of COPD and asthma, it serves as a benchmark for comparing the anti-inflammatory effects of new drug molecules. The concentration 1 × 10^−6^ M caused a two-fold reduction in IL-6 and three-fold reduction in TNF-α concentration, as shown in figure 3.2 and 3.7 respectively. Allergic and non-allergic inflammation can be reduced using Budesonide and *in vitro* studied have shown that it is thousand times more potent than cortisol (Sharafkhaneh, 2010).

As can be seen in figure 3.3 and 3.8 the treatment of LPS-stimulated THP-1 cells with Fenoterol has diminished the concentrations of both cytokines. The *P* value was highly statistic (*P*<0.001) in both cases. NF-κB and Toll-like receptors signalling is inhibited effectively in cells treated with β-2 receptor agonist fenoterol. LPS-induced AMP-activated protein kinase (AMPK) activation releases IL-1β and fenoterol inhibited the cytokine release efficiently. The findings were in concordance with the published research of Wang et.al (2015). Dose-responsive pattern was obtained which confirmed that fenoterol is a standard drug. Comparative study of efficacy can be studied using fenoterol as a standard anti-inflammatory drug.

Our results confirmed that the long-acting muscarinic antagonist (LAMA) Tiotropium is relatively more effective in decreasing the IL-6 and TNF-α expression, than the short-acting muscarinic antagonist (SAMA) Ipratropium, inhaled corticosteroid (ICS) Budesonide and β-2 agonist Fenoterol. The inhibitory potential of Tiotropium is mainly because its low dissociation constant value (Haddad, Mak and Barnes, 1994). Moreover, it has a kinetic selectivity for M_1_ and M_3_ receptors (Barnes, 2000) and due to these reasons Tiotropium has been found to be clinically beneficial for treatment of COPD and asthma (Rice, Kunisaki and Niewoehner, 2007). The inhibition of TNF-α after Tiotropium treatment was voluminous as shown in figure 3.10 and the *P* value obtained was *P*<0.0001.

Ipratropium demonstrated a considerable decline in the cytokine expressions for IL-6 and TNF-α, but statistically the results were not significant as shown in figure 3.4 and 3.9. The experimental data was deemed to be insignificant due to presence of extreme outliers. One to two outliers were observed in the ELISA data which were highly deviated from other observations. A trial statistical test excluding the outlier demonstrated that there was a significant difference between obtained data sets. Irrespective of that, the results for current study were presented without exclusion of any outlier to maintain transparency. Moreover, Ipratropium is an anti-inflammatory and its cytokine reducing properties are distinguished. The source of erroneous observation can be due to increased absorbances of the samples which may be due to excess addition of TMB substrate solution, improper mixing of substrate solution and TMB stop solution or improper drug dilutions of Ipratropium. The outliers have most probably stemmed from imprecise drug dilutions in the investigation of IL-6 and TNF-α levels and from figure 3.4 and 3.9 it is evident that the responses are not in a dose responsive pattern which implies that inappropriate drug dilutions are responsible for such outliers.

### 4.2 Current pharmacotherapies for COPD

Currently there are no appropriate therapeutic agents for the treatment of COPD, it can be only managed by using bronchodilators and inhaled corticosteroids mostly. Bronchodilators such as β-receptor agonists and anti-cholinergic drugs are treatment of choice (GOLD, 2006). Short-acting β-agonist Ipratropium in combination with anti-cholinergic salbutamol effectively reduce the inflammation and exacerbations (Petty, 1995). Moderate COPD is maintained by long-acting beta agonists usually the bronchodilatory effect is maintained for 10-12 hours (Dougherty, Didur and Aboussouan, 2003). The major side effects of this therapy are tachycardia, ventricular arrythmia and hypokalaemia (Tashkin and Cooper, 2004). For management of severe COPD long-acting muscarinic Tiotropium is preferred because of its sustained duration of action (24-36 hours) (Tashkin and Cooper, 2004).

Methylxanthines have the potential to reduce inflammation by various mechanisms such as prostaglandin antagonism and adenosine-receptor antagonism majorly. But due to its does-related toxicities it has been given second-class status. (Barr, 2003). Effective treatments which reduce exacerbations and slowdown disease progression are the unmet needs of COPD patients. The inflammation caused by cytokines and chemokines can be reduced with the help of corticosteroids, but it is ineffective in patients suffering from COPD (Keatings *et al.*, 1997). Keatings *et. al* (1997) confirmed that inhaled corticosteroids show limited clinical benefit. Budesonide reduces IL-6 and TNF- α cytokines are to a limited extent only. Our findings suggest that Budesonide majorly reduces TNF- α expression and it is evident in figure 3.7. The steroid resistance of IL-6 was observed in our results.

### 4.3 Targeting muscarinic receptors for treatment of COPD

Muscarinic receptor antagonists elicit bronchodilatory effect by blocking the cholinergic tone of the airway smooth muscles. They increase the airflow and inhibit mucus secretion. Out of the five muscarinic M_1_, M_2_, M_3_, M_4_ and M_5_ receptor subtypes M_1_, M_2_, and M_3_ receptors subtypes have a critical role in treatment of COPD (Buels and Fryer, 2011). The rationale for targeting muscarinic receptors is that M_3_ receptor alone has the potential to inhibit bronchoconstriction. Furthermore, Ipratropium and Tiotropium specifically act through this receptor. The M_1_, M_2_, and M_3_ expressions in the blood monocytes and sputum of COPD patients are relatively high and inflammation of airways is due to production of leukotrienes B4 (LTB4) (Profita *et al.*, 2005). This indicates that antagonists acting on muscarinic receptor specifically, will be able to reduce the inflammation effectively.

### 4.4 Chemistry and Pharmacology of Ipratropium and Tiotropium

Ipratropium is quaternary ammonium compound, specifically it is N-propyl analogue of atropine as shown in figure 4.1(Ensing *et al.*, 1989). It has poor bio-availability and it not capable of crossing the blood-brain barrier (BBB), this limits its systemic availability and due to this it is highly improbable for Ipratropium to produce any adverse effects. Ipratropium substantially reduces the levels of pro-inflammatory cytokines in bronchoalveolar lavage fluid in rodent models (Buels and Fryer, 2011). In present study ipratropium has demonstrated cytokine inhibition in THP-1 cells but statistically insignificant data was obtained, however the decline in cytokines followed a dose-response pattern. Combination of β-2 agonists with ipratropium is more appropriate treatment to improve lung function (National Asthma Education and Prevention Program, 2007).

**Figure 4.1.**
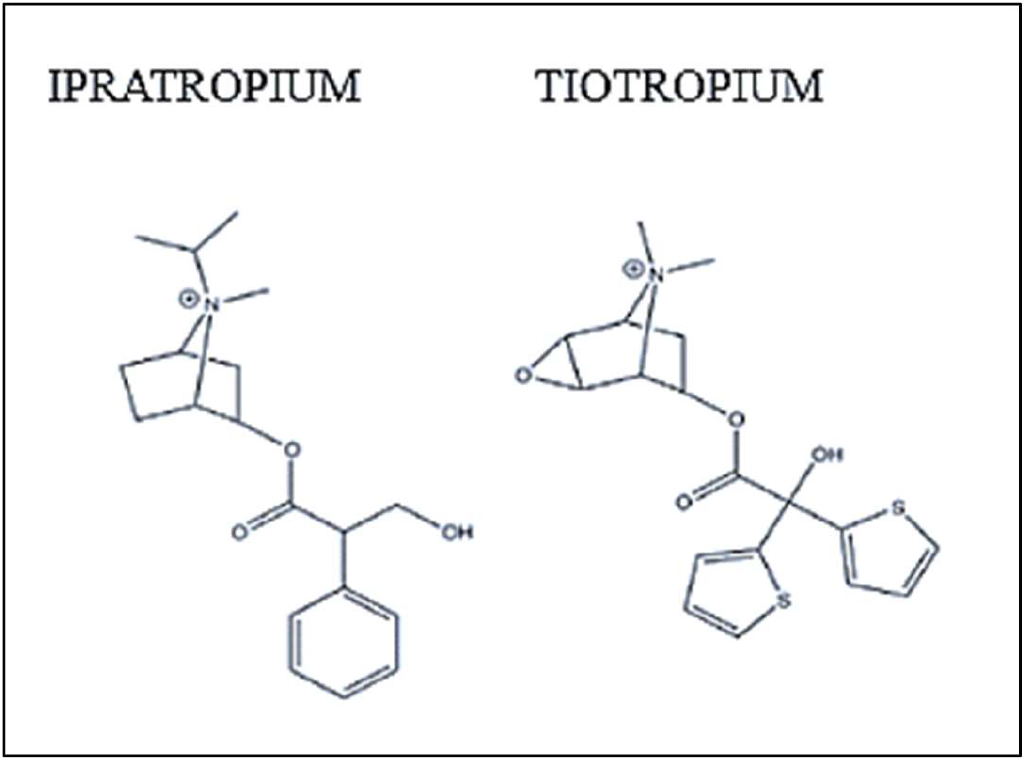
Structure of Ipratropium and Tiotropium.

Ipratropium distinctly blocks the neuronal M_2_ receptors whereas Tiotropium majorly acts through M_3_ receptor and the uptake of these drugs in the airway smooth muscle is mediated by organic cation/ carnitine transporters (OCTN) (Nakamura *et al.*, 2010).

Ipratropium is short-acting mainly because of it dissociates rapidly at the receptor and has low selectivity for M_3_ compared to Tiotropium (Dowling and Charlton, 2009).

Tiotropium has considerably reduced the IL-6 and TNF- α expression in THP-1 cells. The results obtained were significant. However, our data indicates that Budesonide is more effective in reducing the cytokine levels than Tiotropium as shown in figure 3.12. Tiotropium is a novel compound which reduces systemic inflammation effectively and it was first used for the treatment of severe asthma (Peters *et al.*, 2010). Tiotropium is also a quaternary compound derived from atropine and it is structurally identical to Ipratropium. Tiotropium has high specificity and affinity for all the muscarinic receptors (Haddad, Mak and Barnes, 1994).

Tiotropium is a kinetically irreversible M_3_ receptor antagonist and it rapidly dissociates from M_2_ receptors to M_3_ receptors (Moulton and Fryer, 2011). Structural activity relationship studies suggested that the selectivity of M_3_ receptors is due to presence of two thiophene rings in the structure as shown in figure 4.1 and the presence of esterase’s prevents the muscarinic binding of Tiotropium outside the pulmonary airway smooth muscles. Alkyl chain in the chemical structure establishes an additional hydrogen bond with the receptor. The binding of the two heterocyclic rings outside the G-protein coupled receptors (GPCR) active site allows Tiotropium to block the receptor (Hohlfeld *et al.*, 2013).

Structural and dynamic studies of M_3_ receptors revealed that Tiotropium binds to the receptor core and the tyrosine portion surrounds the drug and facilitates hydrophobic interactions of Tiotropium as shown in figure 4.2.

**Figure 4.2.**
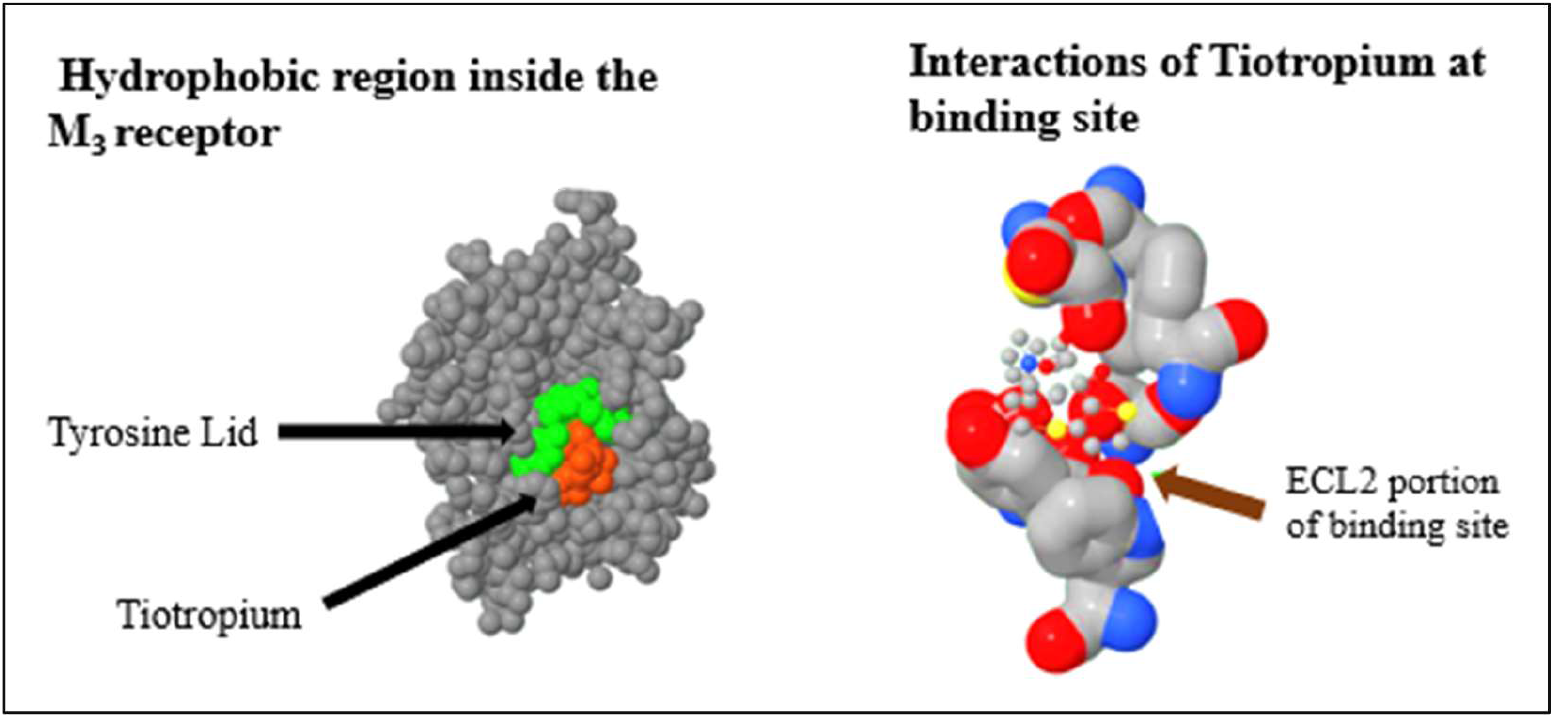
Binding interactions of Tiotropium

The extracellular loop 2 (ECL2) portion of binding site contains Leucine in M_3_ receptor whereas M_2_ receptor has phenylalanine. The substitution of these amino acids creates a large binding site which does not disrupt the hydrogen bonds formed by the thiophene ring present in Tiotropium. These interactions at the receptors are responsible for the slow dissociation which results in sustained duration of action (24-36 hours) (Kruse *et al.*, 2012). The interactions of Tiotropium with the receptor binding site are distinct and these interactions are not exhibited by Ipratropium in M_3_ and M_2_ receptors and presumably this is most reasonable explanation for the sustained action of Tiotropium.

### 4.5 Clinical outcomes of Ipratropium and Tiotropium

Clinical studies involving COPD patients have suggested that Ipratropium significantly improve bronchodilation compared to β-2 agonists Formoterol, but it does not improve the lung function (Montuschi and Ciabattoni, 2015). Similar studies on Tiotropium revealed that Ipratropium is 15 times less potent than Tiotropium. Understanding Potential of Long-term Impacts of Function with Tiotropium [UPLIFT] study revealed that exacerbations and emergency visits to the hospitals were less in patients administering Tiotropium and overall the lung function, and bronchodilation was observed. Whereas the patients administering ICSs and LABA showed effective bronchodilation, but it was not as effective compared to Tiotropium. Additionally, the post-hoc analysis of the study concluded that Tiotropium has least adverse effects (Tashkin *et al.*, 2008). Multiple doses of muscarinic antagonists increase bronchodilation excessively and it can be used to treat moderate and severe COPD (Rodrigo and Castro-Rodriguez, 2005).

Cardiovascular adverse effects associated with the use of Ipratropium (Ogale *et al.*, 2010) and Tiotropium in COPD patients are quite controversial (Singh and Loke, 2010). However, cardiovascular stroke or pneumonia associated risks were not observed in UPLIFT study (Tashkin *et al.*, 2008).

Ipratropium and Tiotropium have been associated with adverse effects such as dryness of mouth, tachycardia and dyspnoea. Ipratropium induces myocardial injury via necrotic and apoptotic mechanisms in animal models (Harvey, Hussain and Maddock, 2014). Tiotropium in combination with LABA/ICSs are associated with cardiovascular events (Liou *et al.*, 2018). Pro-arrhythmic and ischaemic effects are associated with anticholinergics Ipratropium and Tiotropium and more likely to occur in cardiovascular patients (Loke and Singh, 2013).

### 4.6 Research Limitations

The main limitation of this research project is that quantification IL-6 and TNF- α levels was carried out in THP-1 cells but not in human blood, monocytes or peripheral blood mononuclear cells (PBMC). Published research of Schildberger et al critiqued that the effects of LPS on cytokine production in PBMC and THP-1 cells are not identical, overexpression of TNF-α in PBMC and negligible production of IL-6 occurs in such models. (Schildberger *et al.*, 2013). Moreover, interactions of LPS with other cellular components of blood decreases the amplitude of IL-6 and TNF-α expression, which implies that undesirable lipoproteins may appear during *in vivo* analysis and higher concentration of LPS is required to initiate cytokine production. From figure 3.1 and 3.6 it is evident that upon LPS challenge TNF- α is highly expressed than IL-6 and it indicates that a comparative study of test drug and its effect on cytokines will present disproportionate results.

An unrecognized limitation is that only IL-6 and TNF-α biomarkers were measured. Interestingly, the data of COPD patients suggest that the biomarkers C-reactive protein (CRP), IL-8, Surfactant protein-D (SP-D), Chemokine ligand 18 (CC-18) are the biomarkers which are expressed predominantly in the serum of COPD patients (Celli *et al.*, 2012). Instead of measuring the highly expressed biomarkers, the sparsely distributed inflammatory biomarker IL-6 and TNF-α were studied. This implies that Ipratropium and Tiotropium which have shown high suppression of IL-6 and TNF-α should also demonstrate a voluminous effect on reduction of IL-8, CRP, SP-D biomarkers.

Micromolar concentrations 1 × 10^−6^ M, 1 × 10^−7^ M and 1 × 10^−8^ M were utilised during the studies, a more extensive millimolar concentration of drugs should have been tested and proportionally higher LPS concentration should have been used to understand the cytokine reducing effects of the drugs to quantitatively. A meticulous dose-response relationship could have been established using such concentrations.THP-1 cells are immortalized cell lines and they mimic human monocytes closely. Therefore the *in vitro* findings must be interpreted accurately before applying them in animal models and humans. Extensive *in vivo* studies on alveolar macrophages, monocytes, Bronchoalveolar lavage fluid (BALF) and serum are required before confirming the anti-inflammatory properties of the test compounds. The number of replications for the study were only four (N= 4). It is plausible that the inconsistencies with in the data might have influenced the standard means of each group (Vaux, Fidler and Cumming, 2012). As discussed in 4.1 previously, inclusion of one or two extreme outliers might have produced inappropriate results. To ascertain synergistic effects of drugs combinations of SAMA’s, LAMA’s, β-2 agonists and ICS’s should have been used to compare the effectiveness of the Ipratropium and Tiotropium. Despite being highly sensitive and specific ELISA certainly has a few limitations. False positive and negatives data due to insufficient blocking of EIA plate (Leng *et al.*, 2008). The most critical limitation of the study is the type of assay used. Present study was based on published research of Birrell et al (2008). The study was performed using Real Time Quantitative Polymerase Chain Reaction (RT-qPCR). However, in present study ELISA was performed. RT-PCR measures the cytokine mRNA expression. Utilising RT-qPCR could have measured the IL-6 and TNF-α much effectively. This assay is highly sensitive and specific for cytokines and chemokines and there is least probability contamination (Giulietti *et al.*, 2001). The treatment of THP-1 cells with Ipratropium and Tiotropium has elicited cytokine suppression at picogram levels and it is possible that the detection using ELISA was not accurate at such lower concentrations. However, RT-qPCR delivers accurate data for mRNA at initial and exponential phases (Forlenza *et al.*, 2011) and we should employ this technique in further studies.

### 4.7 Future Considerations

Further research utilising various cell lines such as Human mast cell line (HMC-1), Murine monocyte cells, Osteoblast like MG-63 cells can allow us to compare and contrast the cytokine suppression potential of Ipratropium and Tiotropium. Moreover, studies in *in vivo* models using alveolar macrophages, lung tissues, BALF and blood serum are required to allow us to predict the potential of test drugs. The biomarkers of COPD especially IL-8, IL-1β, CRP, CC-18 should be investigated similarly. Investigation of cytokine reduction potential of Salmeterol and Salbutamol (β-2 agonists) with Ipratropium and Tiotropium should be considered in further research. Screening of drug libraries to identify drugs eliciting cytokine suppressing potential in LPS-stimulated cells mimicking the innate immunological inflammatory responses must be studied intensively. The quantification of cytokine production should be conducted using RT-qPCR. Studies which further elucidate mechanisms of IL-6 and TNF-α production and their involvement in COPD are required.

### 4.8 Conclusion

Ipratropium and Tiotropium substantially reduced IL-6 and TNF- α concentrations in THP-1 cells and the research hypothesis was found to be true. Further implications can be drawn that these drugs possess anti-inflammatory potential and in a clinical setting they can be used to reduce chronic inflammation. The results of this research showed that Ipratropium and Tiotropium reduce cytokines, but not as effectively as Budesonide and Fenoterol. It can be assumed that Ipratropium and Tiotropium can markedly reduce inflammation in COPD, and it should be categorised as novel anti-inflammatories for treatment of COPD.

## Notes

### Competing Interest Statement

The authors have declared no competing interest.

